# Proximity labelling identifies pro-migratory endocytic recycling cargo and machinery of the Rab4 and Rab11 families

**DOI:** 10.1101/2022.05.12.491689

**Authors:** Beverley Wilson, Jakub Gemperle, Craig Lawless, Chloe Flett, Matthew Hartshorn, Eleanor Hinde, Thomas Harrison, Megan Chastney, Sarah Taylor, Jennifer Allen, Patrick Caswell

## Abstract

Endocytic recycling controls the return of internalised cargos to the plasma membrane to coordinate their positioning, availability and downstream signalling. The Rab4 and Rab11 small GTPase families regulate distinct recycling routes, broadly classified as fast recycling from early endosomes (Rab4) and slow recycling from perinuclear recycling endosomes (Rab11), and both routes handle a broad range of overlapping cargos to regulate cell behaviour. We adopted a proximity labelling approach, BioID, to identify and compare the protein complexes recruited by Rab4a, Rab11a and Rab25 (a Rab11 family member implicated in cancer aggressiveness), revealing statistically robust protein-protein interaction networks of both new and well characterised cargos and trafficking machinery in migratory cancer cells. Gene ontological analysis of these interconnected networks revealed that these endocytic recycling pathways are intrinsically connected to cell motility and cell adhesion. Using a knock sideways relocalisation approach we were further able to confirm novel links between Rab11/25 and the ESCPE-1 and retromer multiprotein sorting complexes and identify new endocytic recycling machinery associated with Rab4, Rab11 and Rab25 that regulates cancer cell migration in 3D-matrix.

## Introduction

Endocytosis, trafficking through the endocytic system and recycling back to the cell surface are evolutionarily conserved processes that control composition of the plasma membrane, which forms the interface between cells and their microenvironment (Simonetti and Cullen, 2019). Endocytic trafficking regulates the positioning and availability of cell surface receptors, their ability to switch on downstream signalling modules and mediates cell-cell and cell-matrix interactions (Scita and Di Fiore, 2010). Trafficking processes are therefore key to determining cell behaviour in normal physiology, and in the progression of disease.

Endocytic recycling plays key roles across a myriad of cell functions, including nutrient uptake, ion homeostasis, cell plasticity, division, metabolism, polarity and migration in a broad range of model organisms and systems. The Rab and Arf families of small GTPases play critical regulatory roles in endocytosis and transit of cargos through the endolysosomal system (Stenmark, 2009). Rab GTPases act to coordinate trafficking and maintain the fidelity of interactions by recruiting effector proteins that control vesicle/endosome linkage to motor proteins, lipid composition, tethering and fusion with target membranes (Stenmark, 2009). Perhaps the best characterized recycling regulators include members of the Rab4 (Rab4a and Rab4b) and Rab11 (Rab11a, Rab11b and Rab25) families, where Rab4 controls recycling from early endosomes (EEs) and Rab11 controls recycling from recycling endosomes (REs) (Grant and Donaldson, 2009).

Detailed molecular mechanisms coordinated by multiprotein complexes have been revealed to control sorting and fusion events in the endolysosomal pathway. Sorting in EEs is controlled by endosomal sorting complex required for transport (ESCRT)s, which recognize ubiquitinylated cargos destined for degradation, and by specific retrieval pathways that use sorting nexin (SNX)-based recognition of cargo to target them for recycling (Retromer, Retriever/CCC, ESCPE-1; each of which depends on the actin regulatory Wiskott Aldrich Syndrome protein and scar homologue complex (WASH) complex; Simonetti and Cullen, 2019). Fusion regulators include the multi-subunit tethers class C core vacuole/endosome tethering (CORVET; EE:EE fusion), homotypic fusion and vacuole protein sorting (HOPS; late endosomal fusion), class C homologs in endosome–vesicle interaction (CHEVI; transit of cargos through REs), endosome-associated recycling protein (EARP; van der Beek et al., 2019; Schindler et al., 2015), and FERARI (factors for endosome recycling and Rab interactions) is a Rab11 interacting multiprotein complex that is crucial for Rab11 function (Solinger et al., 2020). Emerging evidence is beginning to describe the links between steps in traffic control, for example Rab11 is required for the recruitment of CHEVI to REs (van der Beek et al., 2019), and CHEVI (specifically the Vps33B subunit) can interact with the CCC complex (Hunter et al., 2018), which is in turn presented cargos by retriever (McNally et al., 2017), suggesting that there may be handover between these critical regulatory players.

Recycling of receptor tyrosine kinases (RTKs, including EGFR, c-Met, PDGFR, EphA2), chemokines receptors, and adhesion receptors is implicated in eliciting the signalling and cell-matrix interactions that permit motility in 2D and 3D matrix, contributing to development, immune surveillance and the invasiveness of cancer cells (Wilson et al., 2018). The Rab11 family plays a key role in control of cell migration, cancer cell invasion and metastasis (Paul et al., 2015). Rab25 is upregulated in some cancers, and can increase their aggressiveness when the lysosomal protein CLIC3 is co-upregulated to control a pathway that rescues integrins from lysosomes (Dozynkiewicz et al., 2012). The Rab11 effector protein Rab-coupling protein (RCP; Rab11-family interacting protein (FIP)-1) is similarly implicated in cell migration, invasion and cancer progression (Caswell et al., 2008; Muller et al., 2009; Zhang et al., 2009). In both cases integrins, including α5β1, are key cargoes that drive invasion into extracellular matrix that contains fibronectin (FN), the ligand for α5β1. Rab4-driven recycling has been shown to play a key role in directional migration and cancer cell invasion, by controlling the trafficking of cargoes such as αvβ3 integrin and MT1-MMP (Christoforides et al., 2012; Frittoli et al., 2014), particularly in 3D microenvironments low in FN. Whilst numerous studies have characterized specific GTPases and cargoes that are key to controlling migration and invasion in specific cell contexts, a broad understanding of machineries that control these pro-migratory recycling pathways is lacking.

Here, we have used BioID based proximity labelling to characterize the protein complexes that form around recycling vesicles in a mesenchymal, migratory ovarian cancer cell line model, specifically focusing on Rab4a, Rab11a and Rab25. Our data reveals a protein-protein interaction (PPI) network that overlaps between these three recycling regulators, and links Rab GTPases to previously identified multi-subunit complexes that control trafficking events in the endolysosomal system. In addition, new Rab4a, Rab11a and Rab25 associated proteins are identified that play key roles in the migration of cells in 3D-matrix.

## Results

### Identification of Rab4a, Rab11a and Rab25 interactomes in live cells

Proximity labelling methods are powerful techniques to understand the formation of protein complexes within living cells. We chose BioID (Roux et al., 2012), as the long labelling time is particularly well suited to long-lived or repeated interactions, such as those that occur during cycles of endocytic recycling at steady state. Because Rab GTPase overexpression can lead to mislocalisation (Gemperle et al., 2021), we adopted a system to select for low level expression of BioID-Rab4a, BioID-Rab11a, and BioID-Rab25 (alongside a cytoplasmic BioID control) at a level close to endogenous protein (Figure S1A-C). Rab GTPases localized to endosomal compartments consistent with EEs (Rab4) and REs (concentrated in a perinuclear region; for Rab11 and Rab25; Figure 1A), whereas BioID alone was diffuse in the cytoplasm. Each BioID-Rab efficiently labelled proteins across a range of molecular weights (Figure S1D) and biotinylated proteins co-distributed with BioID-Rabs and were observed at the cell periphery (Figure 1A). We therefore devised a proteomic approach to compare the protein complexes that form around Rab4a, Rab11a and Rab25 recycling vesicles and endosomes, where biotinylated proteins were isolated from cells expressing BioID-Rab4a, BioID-Rab11a and BioID-Rab25 alongside BioID control under stringent conditions using an optimized protocol, before analysis by tandem mass spectrometry and label free quantification (quadruplicate samples, Figure S1E-G).

**Figure 1:**
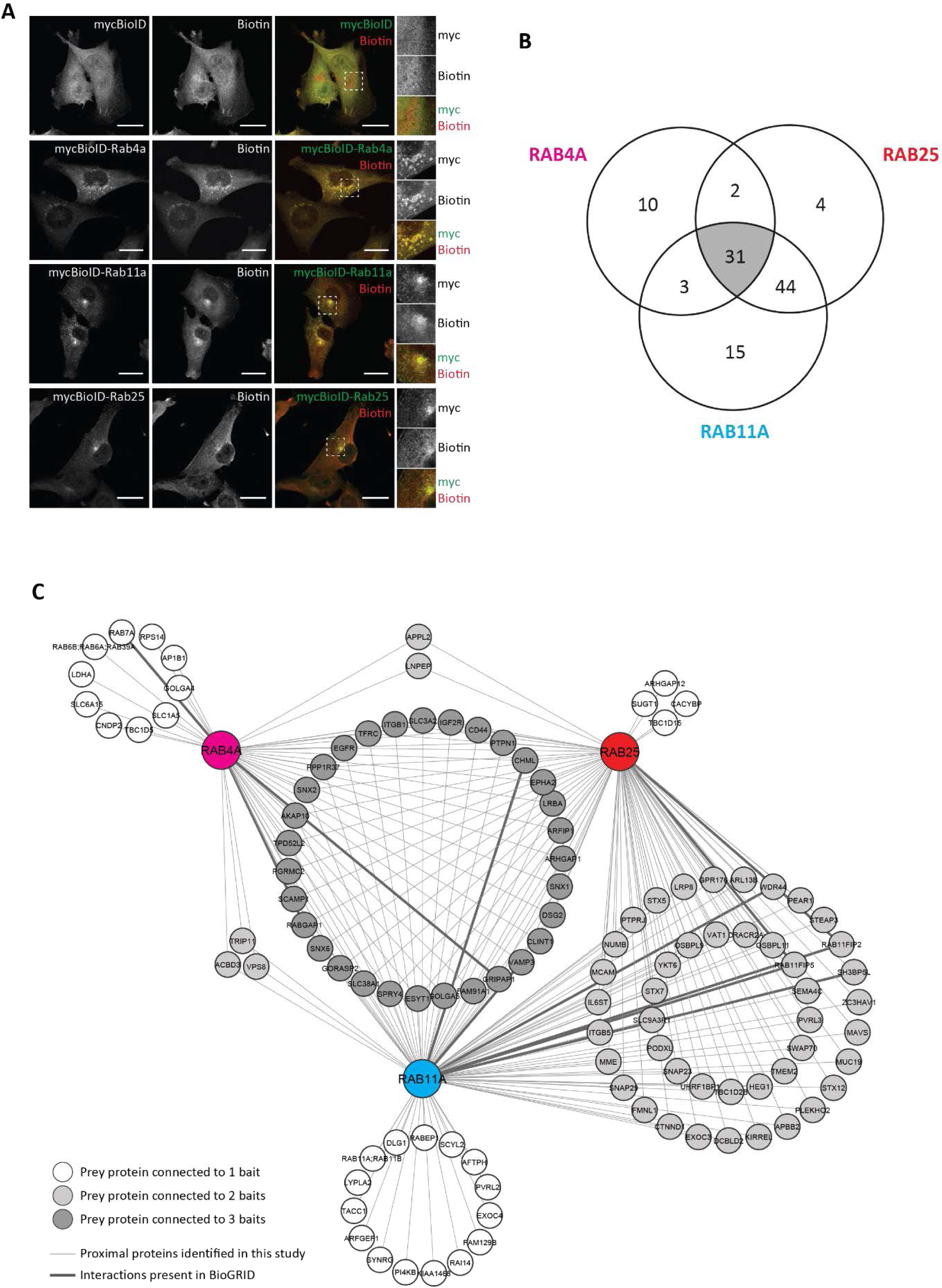
BioID identifies a network of proteins associated by the regulators of endocytic recycling Rab4a, Rab11a and Rab25. **(A)**. Localisation of fusion proteins and biotinylated proteins in A2780 cells stably expressing mycBioID or mycBioID-Rab4a/11a/25 cultured with biotin (1 μM, 16 h). Zoomed regions show the perinuclear region of the cell. Scale bars, 20 μm. **(B)**. Venn diagram showing the overlap of high-confidence proximal proteins (BFDR≤0.05) identified for Rab4a, Rab11a and Rab25 using an optimised BioID protocol (see S1E) that was carried out with four biological replicates (n=4). Samples were analysed by mass spectrometry before data analysis by MaxQuant and SAINT express to identify high-confidence Rab GTPase proximal proteins. **(C)**. Network showing the proteins identified as high-confidence proximal proteins (BFDR≤0.05) for Rab4A, Rab11a and Rab25. Thicker lines connecting proteins to bait Rab GTPases represent interactions found in the BioGRID PPI database. Coloured nodes, bait proteins; white nodes, proteins enriched to one Rab GTPase; pale grey nodes, proteins enriched to two Rab GTPases; dark grey nodes, proteins enriched to all three Rab GTPases.

Comparison of each BioID-Rab with BioID control using robust statistical analysis by SAINT identified a high confidence interactome of proteins (Figure 1B) associated with Rab4a (46), Rab11a (93) and Rab25 (81) which were well distinguished from BioID control by principal component analysis (Figure S1F, Supplementary Table 1). This extended to a ‘longlist’ of 134 proteins associated with Rab4a, 266 with Rab11a and 230 with Rab25 when preys were identified by a fold-change rule (>2-fold enrichment, at least 2 unique peptides, in at least 3 of 4 repeats; Supplementary Table 2). Of the 31 high confidence associated proteins shared by all 3 Rab GTPases, the fold enrichment was closely correlated between Rab11 family members Rab11a and Rab25, but less so with Rab4a (Figure S1G). The high confidence interactomes were significantly enriched for gene ontology (GO) biological processes associated with protein localization and vesicle fusion, although interestingly vesicle docking related terms were only enriched to Rab11a and Rab25 (Figure S2A-C). GO cellular compartments were unsurprisingly dominated by vesicle and endosome related terms, and interestingly cell leading edge, lamellipodium and focal adhesion related terms were enriched for all GTPases (Figure S2A). GO molecular function and biological process terms lamellipodium organisation/morphogenesis were enriched, suggesting that many of the cargoes are involved in processes related to adhesion and protrusion (Figure S2B-C). Rab4a associated proteins were also associated with terms related to amino acid transport (Figure S2B-D), suggesting these metabolic processes could be specifically regulated by this recycling regulator.

### The trafficking machinery associated with Rab4a, Rab11a and Rab25

The Rab associated proteins within our robust proteomic dataset were predominantly a range of cell surface proteins, trafficking machinery, Rab/Arf/Rho regulators and effectors, phosphoinositide regulators and clathrin associated protein (Figure 2). These included many known Rab4a, Rab11a and Rab25 interactors (categorized based on function, Figure 2), for example CHML (CHM Like Rab Escort Protein/Rab escort protein-2) was robustly recruited by all 3 Rab GTPases (Figure 2), but it also revealed specific interactions and previously uncharacterised links to trafficking regulators. Of the well characterized Rab11 family effectors, Rab11FIP5, Rab11FIP2, WDR44, KIAA1468 (RELCH), EXOC3 and OSBP9/11 were robustly recruited to both Rab11 and Rab25 (Figure 2, Supplementary Table 1 and 2) and RCP (Rab11FIP1, longlists only) was also enriched. Interestingly, PI4KB was significantly enriched to Rab11a only, suggesting that whilst many effectors are shared, PI4KB is relatively specific. The intracellular nanovesicle (INV) protein TPD52L2 (TPD54) (Larocque et al., 2020) was recruited to each Rab, but most abundant with Rab11 and Rab25. Rab4a recruited it’s known effectors GRIPAP1 (GRASP1) and RUFY1 (longlist only), although interestingly the Rab effector RABEP1 (Rabaptin-5) was a high confidence Rab11a binding partner that appeared on the lower confidence Rab4 longlist (Figure 2, Supplementary Table 1 and 2). Interestingly, recruitment of other Rab GTPases was suggested by the BioID dataset. Surprisingly, Rab11a was not enriched to Rab25, despite their high sequence similarity. Rab4a strongly recruited Rab6 (Rab6A:Rab6B:Rab39; protein group not distinguishable by MS/MS) and Rab7a, suggesting interplay between these trafficking routes (Figure 2). Furthermore, Rab11a and Rab25 recruited the related GTPases Arl13b (Figure 2), RABL6 (longlist only, Supplementary Table 2) and CRACR2A (Rab46), a dynein adaptor which can play a role in the stimulated release of Weibel-Palade bodies in endothelial cells (Miteva et al., 2019; Pedicini et al., 2021; Wang et al., 2019). Taken together these data give confidence that BioID is an excellent technique to capture and compare Rab interactions in live cells.

**Figure 2:**
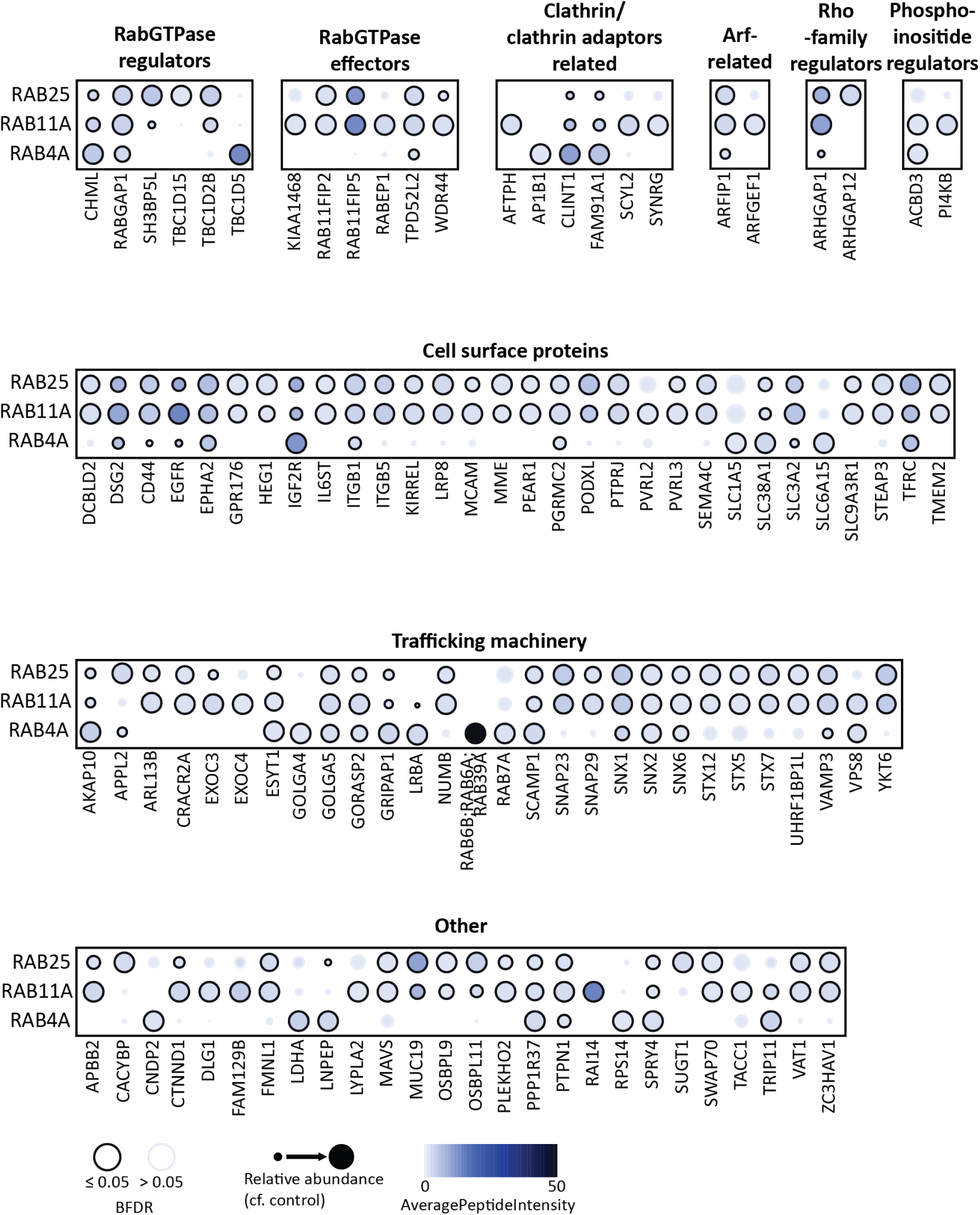
Enrichment of associated proteins to endocytic recycling Rab GTPase baits. Dot plot of the high-confidence proximal proteins was generated using the ProHits-viz online tool (Knight et al., 2017). Proximal proteins are represented by circles and are attributed to each of the three bait Rab GTPases in rows and were manually split into different groups based on their function. Circle size relates to the relative abundance of the protein in that sample (analogous to fold change). Circle colour relates to the average intensity (AveragePeptideIntensity) of the protein in that sample. Circle outline relates to whether the protein is classed as a high confidence enriched protein to that sample (BFDR≤0.05).

#### Guanine nucleotide exchange factors

Whilst GEFs for Rab11 have been identified, for Rab4 these have remained elusive. The TRAPP II complex activates Rab11 homologues in yeast and Drosophila (Lamber et al., 2019; Sakaguchi et al., 2015), but components of this complex were not enriched in our dataset. Recently, SH3BP5 (REI-1) was identified as a mammalian Rab11 GEF, and subsequently SH3BP5 and the closely related SH3BP5L were shown to behave as GEFs for Rab11 and Rab25 in vitro (Jenkins et al., 2018; Sakaguchi et al., 2015). Our data show that SH3BP5L was enriched to both Rab11a and Rab25, albeit with higher levels associated with Rab25 (Figure 2, Supplementary Table 1 and 2). DENND4C, related to Drosophila Rab11 GEF CRAG (Xiong et al., 2012), was enriched to Rab11a (longlist only, Supplementary Table 2), suggesting that this could be an alternate GEF for Rab11. Furthermore, SMCR8 (part of the C9orf72-SMCR8 complex with GEF activity for Rab8 and Rab39) was enriched to Rab25 (longlist only, Supplementary Table 2), which could suggest it has GEF activity specific to this GTPase, but no potential Rab4 GEFs were identified.

#### GTPase activating proteins

GAPs for the Rab4 and Rab11 families are better understood, and of these the Tre-2/Cdc16/Bub2 (TBC) domain family are best characterized. RabGAP1 ((TBC1D11); a Rab2, Rab4, Rab6, Rab11 and Rab36 GAP) was enriched to each Rab tested (Figure 2, Supplementary Table 1 and 2). TBC1D5 (a Rab7 GAP (Müller and Goody, 2018)), was identified as a high confidence Rab4 proximal protein (Figure 2) and TBC1D8 (specificity unknown) was a lower confidence association (longlist only, Supplementary Table 2). AKAP10, the drosophila homologue of which has been proposed to act as a Rab4 GAP and Rab11 GEF (Eggers et al., 2009), was significantly enriched to all Rabs, but most abundant with Rab4 (Figure 2). TBC1D15 (a Rab11a/b GAP (Müller and Goody, 2018)) was identified as proximal to only Rab25, whilst TBC1D2B (a Rab22 GAP (Kanno et al., 2010)), RabGAP1L (TBC1D18; a Rab22a, Rab34 and Rab39b GAP, longlist only), TBC1D5 (longlist only), TBC1D22B (specificity unknown, longlist only) were Rab11 and Rab25 proximal (Figure 2, Supplementary Table 1 and 2). These data highlight the overlapping regulation of Rab GTPases by specific GAPs, but also point to a more complex and interconnected regulation between Rab-specific GEFs and GAPs.

In addition to Rab regulators, GEFs/GAPs for other Ras superfamily members were also identified (Figure 2; Supplementary Table 1 and 2), predominantly for Arf and Rho families, perhaps pointing towards the influence of vesicle trafficking on cytoskeletal signalling (Jacquemet et al., 2013; Wu et al., 2011) and Rab-Arf interconnected cascades (D’Souza et al., 2014). RhoGTPase regulators included ARHGAP1 (p50RhoGAP) previously identified as a vesicular protein (Sirokmány et al., 2006), enriched to all Rabs but most abundant with Rab11 and Rab25, ARHGAP12 (Rab25 only), ARHGAP18 (Rab11 longlist only) and ARHGEF1 (p115RhoGEF, Rab11 longlist only). Arf regulators included ARFGEF1 (BIG1, Rab11 only), ARFGEF2 (BIG2, Rab4 longlist only), ARFGAP1 (Rab11 and Rab25 longlist only) and ARFGAP3 (all Rab longlists only).

#### Multiprotein trafficking complexes

Rab11FIP5 of the FERARI complex was highly enriched to Rab11 and Rab25 (but not Rab4; Figure 2), and FERARI complex components EHD1 (Rab11 and Rab25 longlists only) and VPS45 (Rab11 longlist only) were also identified, confirming the ability of BioID to identify trafficking complexes. Interestingly, VPS33B (CHEVI complex) was recruited by Rab11 (longlist only), and VPS8 was identified with all Rabs (Figure 2; Rab25 longlist only Supplementary Table 2). VPS51 (Ang2), a component of the endosomal associated recycling protein (EARP) tether complex previously seen to localise to Rab4 and Rab11 endosomes (Schindler et al., 2015) was enriched to all Rabs (longlists only) together with interaction partner tSNARE syntaxin 6 (STX6; longlists only). Interestingly, RETRIEVER and COMMD complex components were not identified, but SNX1, SNX2, and SNX6 were robustly identified with all Rabs (Figure 2; Supplementary Table 1 and 2). SNX3 (longlists only) also associated with all Rabs, whilst SNX4 (Rab11 longlist), SNX29 (Rab4 longlist), FAM21A (WASH complex component, Rab11 and Rab25 longlist) were more selectively recruited. These data point towards a previously unknown association between Rab4, -11, and -25 recycling machinery and sorting nexins associated with the ESCPE-1 (SNX1/2/6) and retromer (SNX3) recycling complexes.

### Detection of directly biotinylated peptides reveals proximal domains between interactors

BioID involves direct biotinylation of lysine residues in close proximity to the bait protein, which can be clearly detected as a modification by mass spectrometry. For the majority of proteins, the biotin modification was not specifically identified on a peptide, however for any peptides it was identified on this would indicate a more significant higher affinity, longer term or repeated interaction (Figure S3A). This analysis revealed the overlapping interaction domains involved between associated trafficking machinery/cargoes and Rab4, -11 and -25, but also highlighted key differences in proximal proteins (Fig 3A-C, Figure S3B-H, Supplementary Table 3).

**Figure 3:**
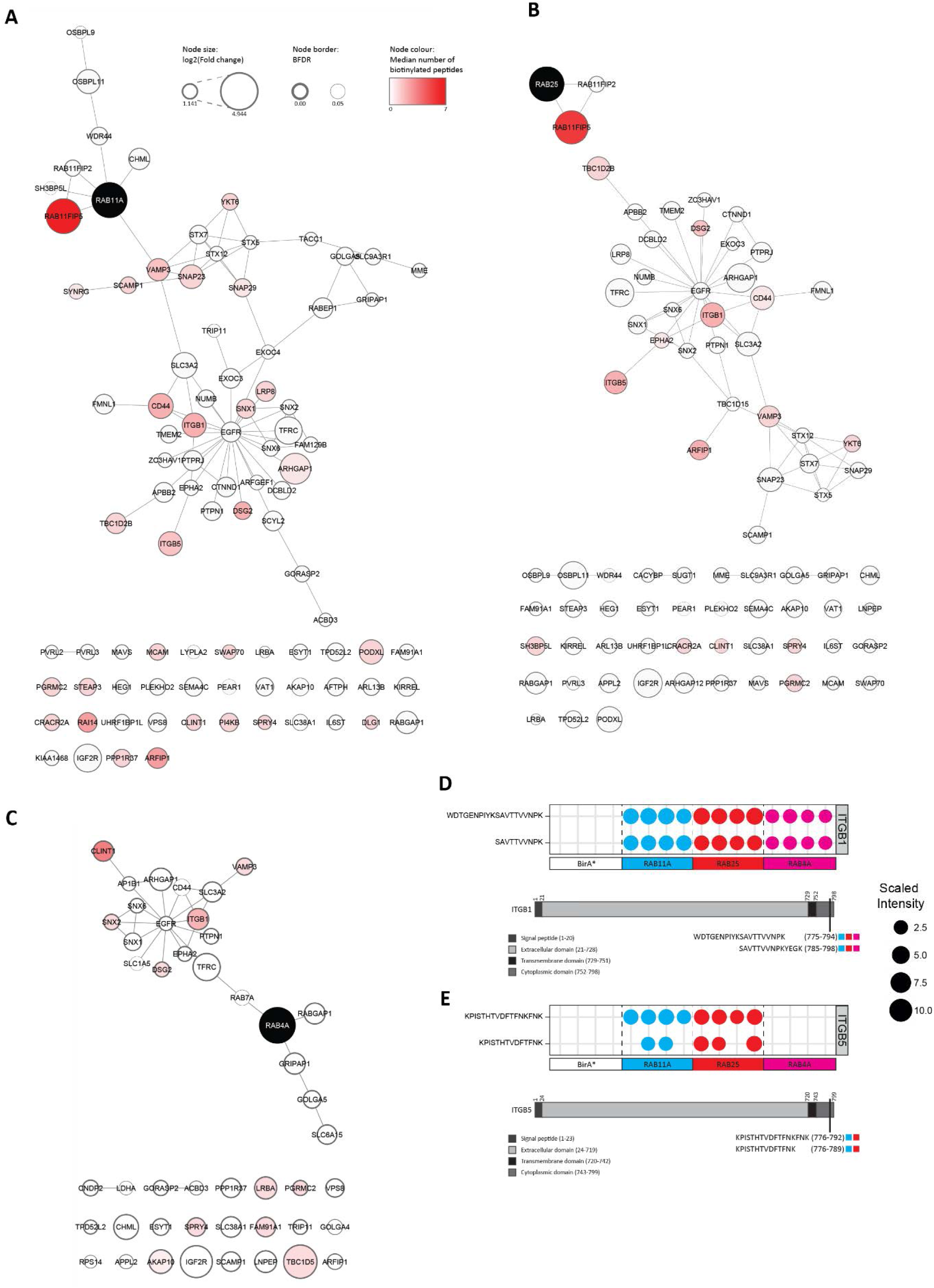
Biotin modified peptides reveal proximal interactors within PPI networks. High-confidence Rab4a **(A)**, Rab11a **(B)** and Rab25 **(C)** proximal proteins (Figure 1C) were mapped onto a PPI database. Rab GTPase nodes coloured black, other nodes are the high-confidence proximal proteins identified for each Rab GTPase. Edges represent interactions present within the PPI database. Node size, log2fold change; node border, BFDR from SAINTexpress analysis; node colour, the median number of biotinylated peptides identified. Biotinylated integrin beta-1 **(D)** or beta-5 **(E)** peptides were mapped to protein domains. Protein domains are shown by grey colourings, domains are not to scale. The position of the biotinylated peptides in the protein are indicated, and coloured boxes show in which samples they were identified; Rab11a, blue; Rab25, red; and Rab4a, pink.

Rab11FIP5 is a key effector of Rab11 family GTPases, and was heavily biotinylated by both Rab11a and Rab25 but not by Rab4a (Figure 3A-C, S3C) in a region within 200 residues of the C-terminal Rab-binding domain, but away from the N-terminal C2 domain, consistent with the known site of interaction (Prekeris et al., 2001). Cargoes such as ITGB1 (β1-integrin) were detected in similar abundance between Rabs, and showed a remarkably consistent level of biotinylation in 2 peptides that overlap (Figure 3D) within the cytoplasmic domain of the integrin (Figure S3B) that was previously found to interact with Rab25 (Caswell et al., 2007). ITGB5 was detected as a biotinylated cargo for only Rab11 and Rab25 (Figure 3E, S3B), suggesting that this integrin may be more selectively recycled via Rab11 family members. In both cases biotinylation pinpointed the location of proximity adjacent to/within the membrane distal NxxY motif (Figure S3B) thought to interact with the recycling regulator SNX17 and adhesion complex protein kindlin (Böttcher et al., 2012; Steinberg et al., 2012), neither of which were biotinylated in our dataset (although kindlin-2 (FERMT2) was in the Rab11a longlist (Supplementary table 2)).

Rab4 showed selective biotinylation of FAM91A1, previously identified on Golgi-recruited to endosomal vesicles ((Shin et al., 2017); Figure 3C). Furthermore, whilst CLINT1 (clathrin interactor 1/epsinR,) was identified as a Rab4, -11 and -25 proximal protein (Figure 2), it was more heavily biotinylated close to the cargo selective N-terminal ENTH domain by Rab4 (Figure 3C, Figure S3D). CLINT1 interacts with the clathrin adaptor AP1 to mediate TGN-lysosome trafficking of cathepsin D and localises to early endosomes to control retrograde trafficking (Mills et al., 2003), but its function with regard to Rab4 is not clear.

PI4KB, a phosphoinositide-4 kinase, was selectively biotinylated by Rab11a (Figure 3A, S3E) suggesting a specific involvement on Rab11a mediated processes. Interestingly, PI4KB plays a critical role in recruiting Rab11 and its effectors to Golgi membranes (Graaf et al., 2004), perhaps suggesting the interaction and function on this organelle is not shared with the closely related Rab25. Similarly, VPS51 was found to be selectively biotinylated by Rab25 (Figure S3F), which could indicate a more close or persistent interaction of Rab25 with the EARP complex. Both Rab11a and Rab25 (but not Rab4) identified CRACR2A as a potential interactor (Figure S3G). Interestingly, of the potential GEFs, only SH3BP5L was biotinylated, and only by Rab25. This biotinylation occurred within the GEF domain and could suggest SH3BP5L is a more dominant Rab25 GEF, and/or that the interaction is repeated/more stable (Figure 3B, S3H). Of the putative GAPs, only TBC1D5 was biotinylated by Rab4, and only TBC1D2B by Rab11 and Rab25 (Figure 3A-C, Supplementary table 3). SNARE proteins showed a varying degree of selective proximity: YKT6 and VAMP3 were biotinylated by all Rabs, VAMP2 appeared more proximal to Rab11a and Rab25, and SNAP23 and SNAP29 were most heavily biotinylated by Rab11a (Figure 3A-C, Supplementary table 3). Taken together, these data indicate that whilst the abundance of prey proteins in BioID experiments can be extremely informative, close analysis of biotinylated peptides and residues provides a further level of granularity that could be important in deducing functional interactions.

### Knock sideways of bait can validate high affinity prey interactions

Conventional fluorescence microscopy provides good insight into the co-distribution of proteins within cells and is valuable as a tool to validate the types of interactions detected by BioID that are often missed by conventional pulldown strategies. However, co-localisation analysis is insufficient to provide information on relative strengths of interactions within cells. We therefore adapted the knock-sideways approach (Robinson et al., 2010), whereby a mitochondrially targeted-FRB is co-expressed with FKBP fused to a ‘bait’ protein of interest, and rapamycin treatment triggers binding of FKBP to FRB and mitochondrial targeting of the bait, reasoning that strong/repeated prey interactors would be more likely to follow their ‘bait’ upon relocalisation to mitochondria (Figure 4A). We first focused on Rab11-FIP5, as this is a known high affinity interactor of (active) Rab11a and Rab25, but not Rab4. Co-expression of FRB-mito with FKBP-GFP-Rabs and mCherry-Rab11-FIP5 revealed that Rab proteins quickly redistributed to mitochondria upon addition of rapamycin (Rab4: T_1/2_ = 2.3 min ±0.4; Rab11: T_1/2_ = 4.3 min ±0.5; Rab25: T_1/2_ = 5.4 min ±0.1;), whereas mCherry-FIP5 redistributed to mitochondria over a slower timescale (T_1/2_ = 13-26 mins) but only when co-expressed with FKBP-GFP-Rab11a (T_1/2_ = 13.88 min ± 1) or -Rab25 ((T_1/2_ = 24.3 min ± 2.3; Figure 4B, C, S4A). When relocalisation of endogenous Rab11-FIP5 was assessed in knock sideways experiments, relocalisation was again only observed in the presence of rapamycin when FKBP-Rab11a and -Rab25 were expressed (Figure S4B-D). We next tested a second known interactor, PI4KB, which our data indicated may specifically associate with Rab11a, but not Rab4a or Rab25. Indeed, whilst FKBP-GFP-Rab11a redistributed endogenous PI4KB to mitochondria in the presence of rapamycin to a significant extent, Rab4a and Rab25 were inefficient in this regard (Figure S4E). These data indicate that knock sideways of bait proteins can act as a tool to validate potential interactions identified by BioID and strengthen the notion that PI4KB is a more robust interactor of Rab11a than of Rab25.

**Figure 4:**
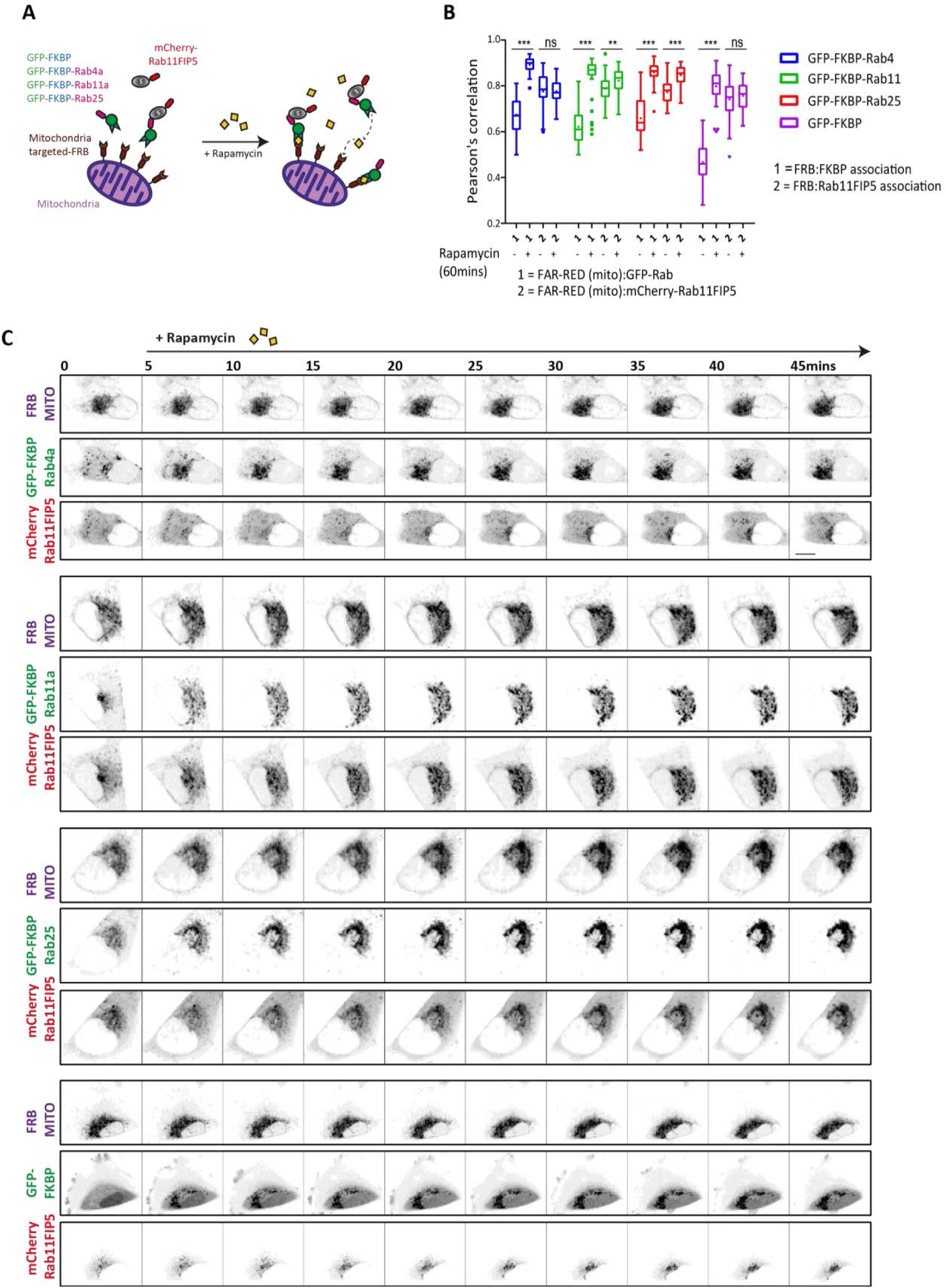
Knock-sideways validation of proximity labelling-identified preys. **(A)**. Schematic illustration of knock-sideways, where mitochondria-targeted FRB/rapamycin is used to induce the re-localisation of GFP-FKBP-tagged Rabs and associated protein complexes. **(B, C)**. A2780 cells expressing iRFP670-FRB and GFP-FKBP or GFP-FKBP-Rab4a/11a/25 and mCherry-Rab11-FIP5 were imaged by spinning disk confocal microscopy before and after the addition of rapamycin (200 nM) to visualise the redistribution of GFP-tagged protein and corresponding localisation of mCherry-Rab11-FIP5. Re-distribution of GFP-FKBP fusion (1) and mCherry-Rab11-FIP5 (2) were analysed by Pearson’s correlation (B; at least 30 cells/condition; statistical analysis with ANOVA/Holm-Sidak post-hoc test; **p<0.01; ***p<0.001), representative images from at least 3 independent timelapse experiments are show in (C; scale bar=10μm).

We next tested knock-sideways efficiency in the relocalisation of sorting nexins that were strong hits in BioID abundance, but more variably biotinylated. SNX1 was identified by Rab4a, Rab11a and Rab25 (Figure 2), but only showed relatively consistent biotinylation with Rab11a (Figure 3A-C). Consistent with this, GFP-FKBP-Rab11a significant redistributed endogenous SNX1 to mitochondria in the presence of rapamycin, but Rab4a and Rab25 knock sideways did not induce mitochondrial localisation (Figure 5A). SNX2 showed similar abundance across Rabs, but SNX2 was only rerouted to mitochondria in cells expressing FKBP-GFP-Rab11a and -Rab25 (Figure 5A, S5A). SNX3 was identified by all Rabs (longlist only), but only significantly redistributed in FKBP-GFP-Rab25 expressing cells (Figure 5A, S5B). Endogenous SH3BP5L (Figure 5B) and CRACR2A (Figure 5C) were significantly redistributed to mitochondria by FKBP-GFP-11a and -25, but not -Rab4a. This is consistent with BioID data (Figure 2), however, biotinylated SH3BP5L peptides were only identified for Rab25 while biotinylated CRACR2A peptides were observed for both Rab11a and Rab25 (Figure S3G-H). Our Rab4 BioID network showed enrichment of CLINT1 and its interactor AP1B (Figure 1C, Figure 2), and CLINT1 was itself directly biotinylated in Rab4 samples (Figure 3C, S3D), however knock-sideways experiments showed that Rab4a was unable to induce significant re-localisation of CLINT1 (Figure S5C), despite good co-localisation between Rab4 and CLINT1 (Figure S5D). Together, these data confirm key findings of proximity labelling but also point towards a more complex relationship between BioID abundance, direct detection of biotinylation, the affinity of bait for prey and the ability to redistribute potential prey proteins.

**Figure 5:**
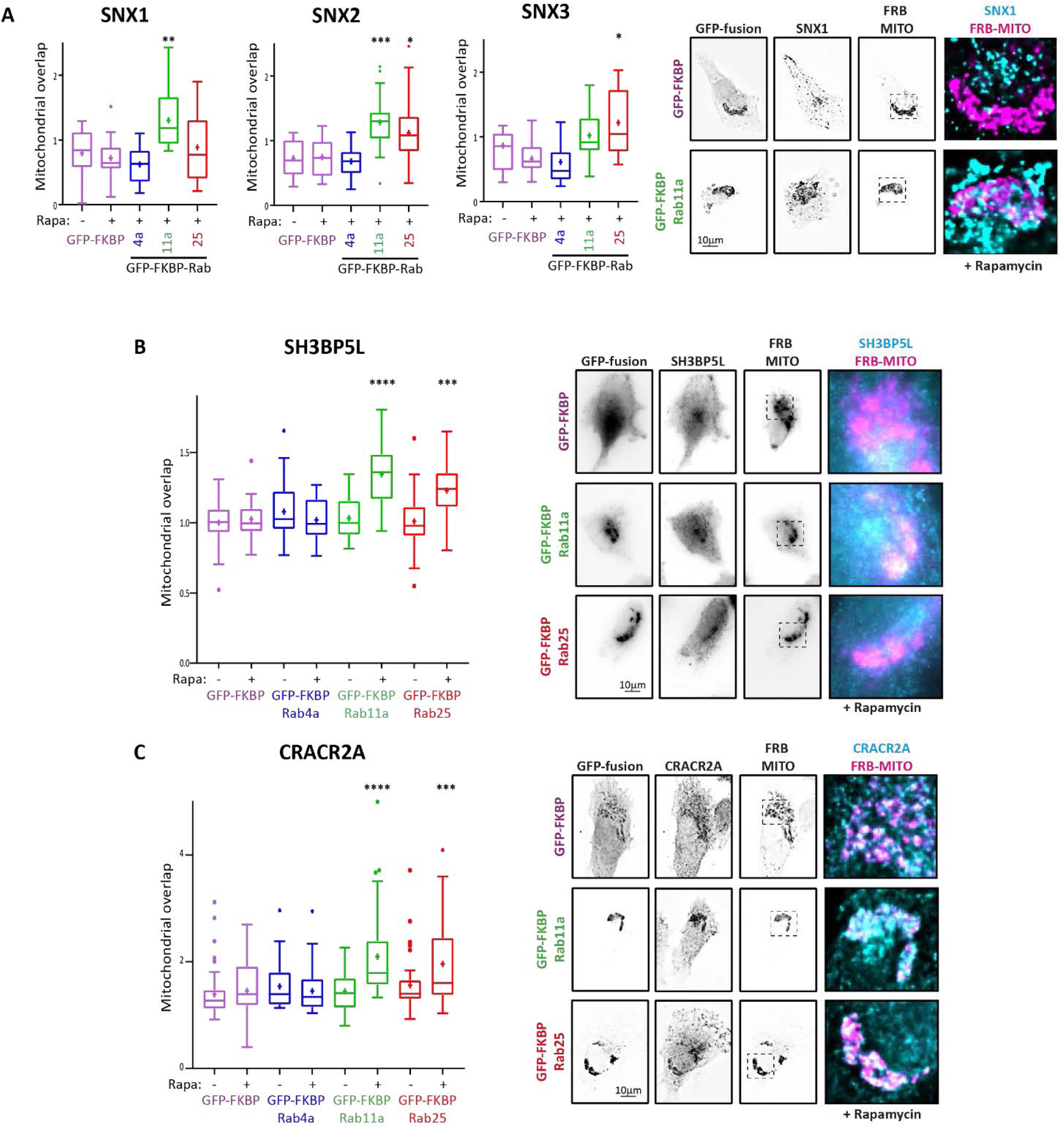
Trafficking machineries are selectively recruited to recycling Rabs. A2780 cells expressing iRFP670-FRB and GFP-FKBP-Rabs treated with rapamycin (200 nM 4h) were fixed and stained for endogenous SNX1/2/3 **(A)**, SH3BP5L **(B)**, or CRACR2A **(C)**. Redistribution of candidate trafficking machinery was analysed by quantifying mitochondrial overlap (see methods; 9-47/cells condition; statistical analysis indicated compared to GFP-FKBP +rapamycin control using Kruskal-Wallis test with Dunn’s multiple comparison; *p<0.05, **p<0.01, ***p<0.01, ****p<0.0001). Representative images from at least 3 independent experiments are shown (scale bar=10μm).

### CLINT1, CRACR2A and SH3BP5L are linked to cell migration within 3D cell-derived matrix

Endocytic recycling pathways regulated by Rab4, Rab11 and Rab25 have each been implicated in cell migration and invasion in 3D matrix. Rab4 controls Rac-driven lamellipodial migration of cells in 2D and invasion in low-fibronectin (FN) through the recycling of αvβ3, whereas Rab11 and Rab25 control α5β1 trafficking to influence migration in high-FN 3D matrix (Caswell et al., 2007, 2008; Christoforides et al., 2012; Dozynkiewicz et al., 2012). We therefore selected Rab4/11/25 associated proteins from our dataset that had not previously been implicated in cell migration/invasion for further analysis in cells that lack Rab25 (A2780-DNA3 where migration is supported by Rab4 (Christoforides et al., 2012) and Rab11 (Gemperle et al., 2021), or express Rab25 at a level similar to those found in ovarian cancer (Cheng et al., 2004) (A2780-Rab25) and migrate in a Rab25/α5β1-dependent manner (Caswell et al., 2007; Dozynkiewicz et al., 2012).

Because CLINT1 was enriched, directly biotinylated in Rab4 samples and co-localized with Rab4 (Figure 3C, S3D, S5D), we focused on the role that CLINT1 plays in cell migration in 3D-cell derived matrix (CDM). Knockdown of CLINT1 with either of two siRNA oligos significantly reduced cell migration in 3D-CDM (Figure S6A, B, E), but only in cells that lack Rab25. This is consistent with the notion the CLINT1 plays a role in Rab4-(but not Rab25) dependent migration in 3D-CDM. We next investigated a Rab11a-specific candidate, PI4KB, which was enriched to and directly biotinylated in Rab11 samples (Figure2, 3A, 3B, S3E), and could be significantly re-localised by knock-sideways of Rab11a alone (Figure S4D). Knockdown of PI4KB, however, did not impact on migration (Figure S6C, E).

SH3BP5L is an interesting candidate shown to behave as a Rab11a/Rab25-specific GEF in vitro (Jenkins et al., 2018), and our data indicate SH3BP5L is significantly enriched to Rab11a but to a greater extent with Rab25 (Figure 2), and that whilst both Rab11a and Rab25 knock-sideways relocalised SH3BP5L (Figure 5B), biotinylated SH3BP5L peptides only appear in Rab25 samples (Figure 3B, S3H). SH3BP5L knockdown had no effect on 3D migration in cells lacking Rab25, but significantly decreased migration of cells migrating in a Rab25-dependent manner (Figure 6A, S6E). These data suggest that SH3BP5L could be specifically required to maintain Rab25 activity in cells migrating in 3D matrix.

**Figure 6:**
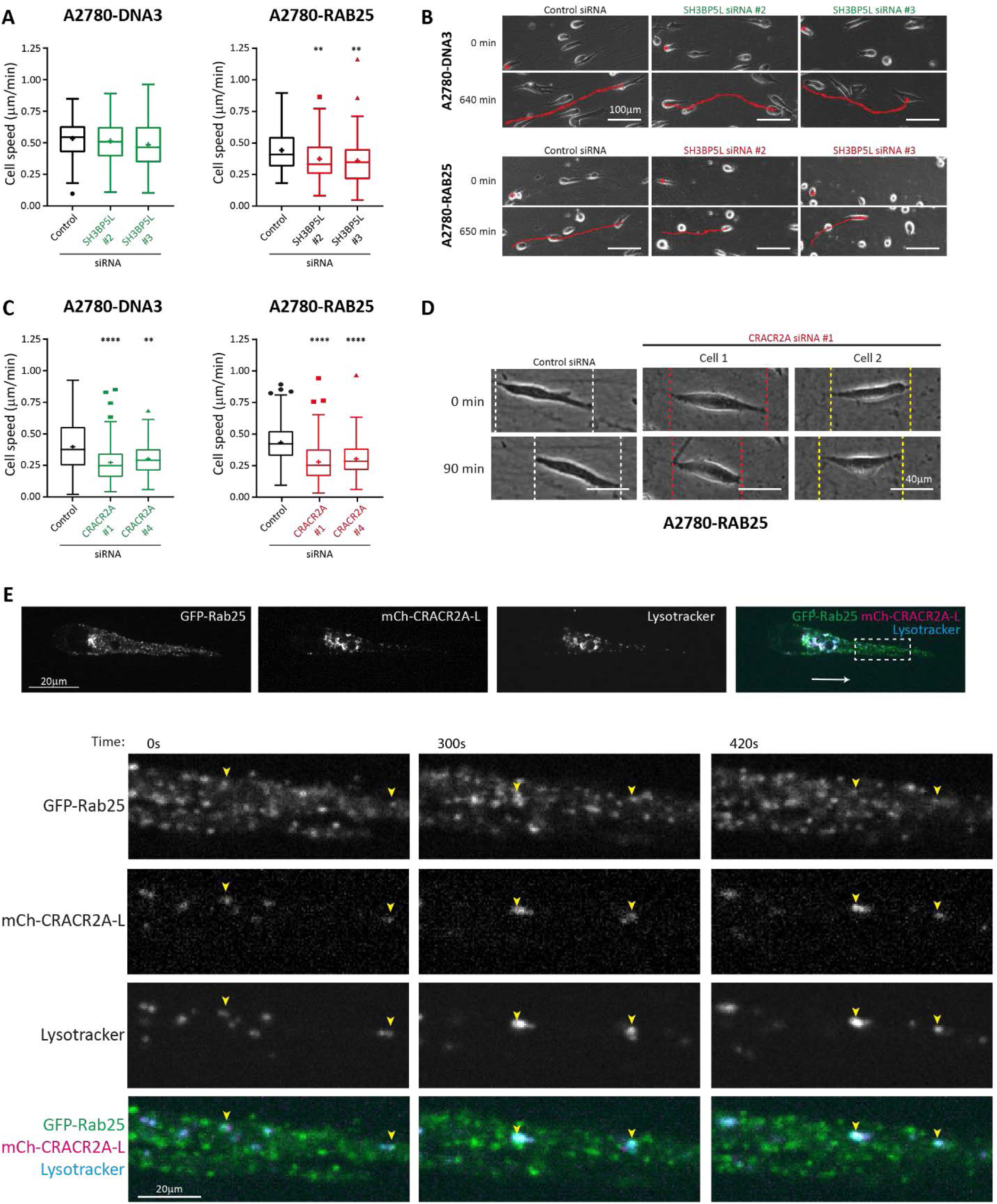
SH3BP5L and CRACR2A are required for cell migration in 3D-matrix. (A, B). A2780-DNA3 and A2780-Rab25 cells were depleted of SH3BP5L by siRNA and seeded into cell-derived matrix for 4 hours before migration was analysed by brightfield timelapse imaging for >16 hours. Cell speed was analysed by manual tracking (A; n≥90 cells from 3 independent experiments; statistical analysis with Kruskal-Wallis/Dunn’s multiple comparisons test; **p<0.01); representative images from at least 3 independent experiments are shown (D; scale bars=100μm). (C, D). A2780-DNA3 and A2780-Rab25 cells were depleted of CRACR2A by siRNA and migration analysis performed as above. Cell speed was analysed by manual tracking (C; n≥90 cells from 3 independent experiments; statistical analysis with Kruskal-Wallis/Dunn’s multiple comparisons test; **p<0.01, ****p<0.0001); representative images from at least 3 independent experiments are shown (D; scale bars=40μm). (E). A2780 cells expressing GFP-Rab25 and mCherry-CRACR2A were seeded into cell-derived matrix for 4 hours before time-lapse imaging by spinning disk confocal microscopy. Dashed white box indicates area enlarged in lower panels. White arrow indicates direction of migration; yellow arrowhead indicates position of CRACR2A positive late endosomes/lysosomes that show overlap with Rab25. Images are representative of 3 independent experiments (scale bars=20μm).

Our data suggest that CRACR2A association is highly significant to both Rab11a and Rab25 to almost identical extents (Figure 1C, 2, 3A, 3B, S3G, 5C). Interestingly, CRACR2A knockdown significantly suppressed migration in 3D-CDM in both Rab25 expressing and non-expressing cells (Figure 6C, S6D, S6E). In both cell lines, CRACR2A depleted cell lines showed a ‘stalling’ phenotype (Figure 6D, S6D), reminiscent of Rab25-expressing cells that lack the lysosomal protein CLIC3 (Dozynkiewicz et al., 2012). mCherry-tagged CRACR2A localised to lysosomes within the cell body (similar to CLIC3 distribution (Dozynkiewicz et al., 2012)), and to lysotracker-positive vesicles, which also contained Rab25 towards the leading edge of pseudopodial protrusions generated in CDM (Figure 6 E). These data suggest that the dynein adaptor CRACR2A is required for efficient migration and may associate with Rab25 to deliver cargoes for trafficking to lysosomes.

## Discussion

Here we show that BioID can be deployed to comparatively analyse the interactomes of Rab-family GTPases and reveal new trafficking machinery that controls cell motility in 3D-matrix. This generated a robust catalogue of Rab4a-, Rab11a-, and Rab25-associated proteins within migratory A2780 ovarian cancer cells, providing an important resource for the further understanding of how trafficking networks perform their myriad function-including in cell migration (Figure 7).

**Figure 7:**
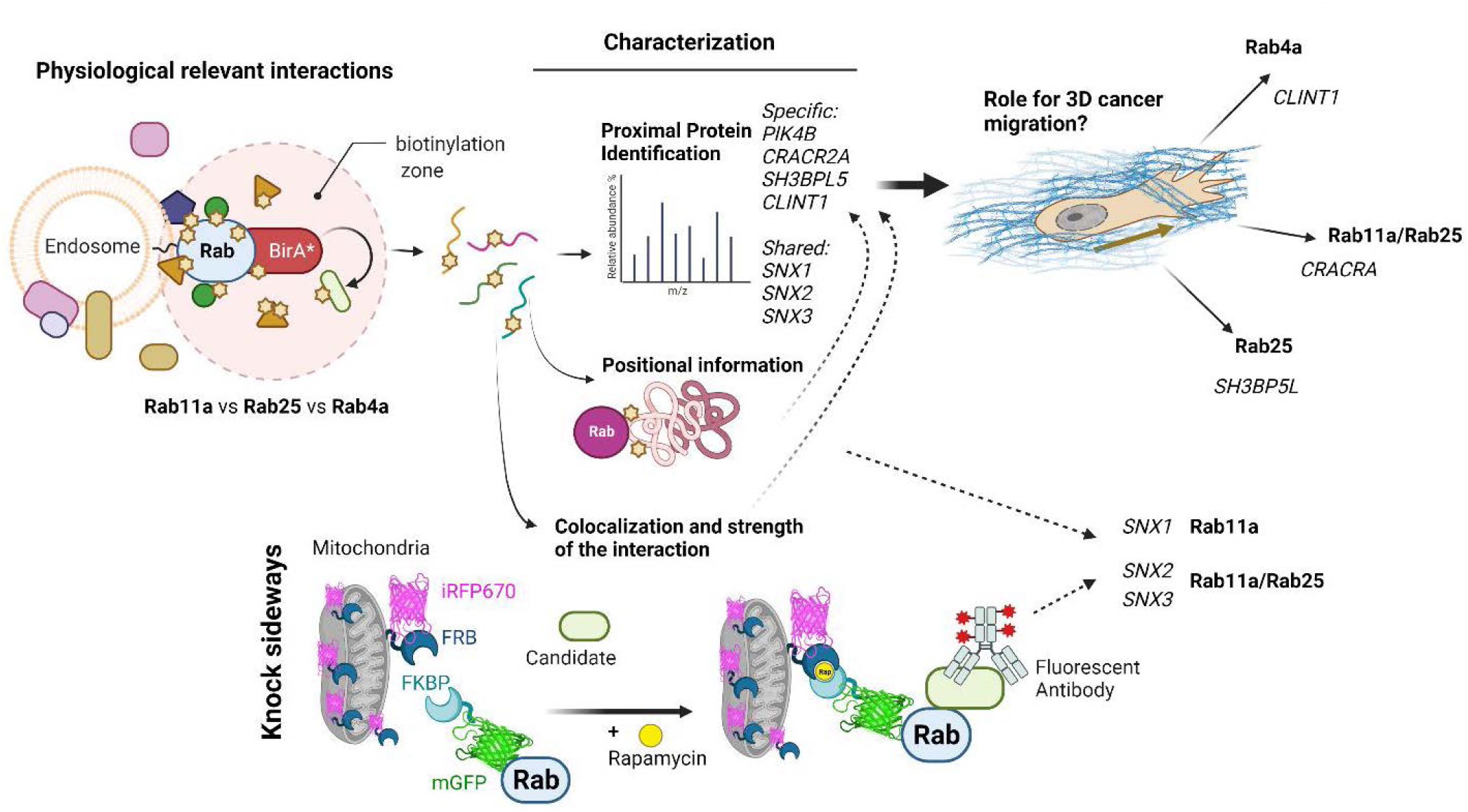
Identification of trafficking machinery and cargoes that promote cancer cell invasion. We describe a pipeline for the identification of cargoes and trafficking machinery associated with Rab4a, Rab11a and Rab25 in migratory cancer cells, coupled with validation of interactions and/or involvement in invasive migration. Schematic illustration was created with BioRender.com

Proximity labelling methods have proven an excellent tool for identification of protein complexes, including for Rab4 and Rab11 GTPases (Gillingham et al., 2019; Go et al., 2021; Liu et al., 2018; Del Olmo et al., 2019). Two studies include both Rab4a and Rab11a in their analyses; Go et al. use BioID to map the proteome in HEK-293 cells (spectral counting based quantification, 2 replicates) (Go et al., 2021), whereas Liu et al. combine affinity purification and BioID in a ‘multiple approaches combined’ (MAC)-tagging strategy (HEK-293 and cancer cell lines, spectral counting based quantification, 2 replicates) (Liu et al., 2018), but neither allow for direct quantitative comparison between GTPases. MitoID combines BioID with mitochondrial tagging to provide a standardised subcellular localisation, and was used to identify potential interaction partners of active and inactive Rab11 mutants in HeLa cells (spectral counting based quantification, 3 replicates) (Gillingham et al., 2019), and an APEX2-Rab4a dataset was also obtained in HeLa cells (spectral counting based quantification, 3 replicates) (Del Olmo et al., 2019). Our data show good overlap with these studies but differs primarily in the direct comparison between Rab4a, Rab11a and Rab25 analysed using quantitative label free proteomics to generate a statistically robust interaction network linking these three recycling regulators. We also further analysed our data by using directly biotinylated peptides to provide another level of confidence, for example the association between Rab4a and Rab11a/25 with the GAPs TBC1D5 and TBC1D2B respectively. Furthermore, we report the first description of protein interaction networks formed by the cancer-relevant Rab25.

Rab11a and Rab25 share 66 % sequence identity at the protein level, differing primarily in their membrane targeting C-termini. It is therefore unsurprising that significant overlap is seen within the cohorts of associated proteins, with particularly high correlation between the level of enrichment to Rab11a/Rab25 for the 31 proteins recruited to Rab4a, Rab11a and Rab25 (whereas recruitment to Rab4a was less well correlated; Figure S1G)). In light of this it is therefore surprising that Rab11a was not found to associate with Rab25, even though there is an indication of Rab4 association with both Rab11a and Rab25. Dimeric Rab11-FIP structures have been described to recruit two copies of Rab11 to form tetramer, but our data hint that these may be homotypic in nature such that Rab25 and Rab11a may function separately. GO analysis indicates that, at least in the migratory cancer cells used in these experiments, the primary functions of these recycling Rabs relates to vesicle and endosome function, but also to focal adhesion and leading edge related processes suggesting broad links to adhesion/migration (Figure S2). In addition, amino acid transport is a significantly enriched GO term for Rab4a associated proteins (Figure S2), indicating that transporter trafficking may constitute an important role of Rab4a that is relatively understudied.

### New trafficking machinery linked to Rab4a, Rab11a and Rab25

In addition to well characterised trafficking machinery previously implicated in Rab4/11/25 recycling, our datasets also provide novel links between the recycling Rabs and other proteins/complexes implicated in different aspects of vesicle trafficking, and new twists on known interactions. For example, the EARP complex component VPS51 is enriched to all Rab’s tested (Supplementary table 2), but only biotinylated directly by Rab25 (Figure S3F), which could suggest a more direct role for EARP in Rab25-mediated trafficking events. Sorting nexins such as SNX4 (identified here in the Rab11 longlist) have previously been associated with Rab11-dependent recycling (van Weering et al., 2012)(Solis et al., 2013), but our experiments also identified ESCPE-1 components (specifically SNX1/2/6) and the retromer component SNX3 (Figure 1C, 2, 5A, S5A-B). Both ESCPE-1 and retromer function in tandem with the WASH complex, of which FAM21A was detected for Rab11a and Rab25. In fact, SNX1 was biotinylated in 3 of 4 Rab11 samples and selectively re-localised by Rab11a knock sideways. Both Rab11a and Rab25 were able to re-localise SNX2, whereas Rab4 was not able to do so. This points towards a closer association between ESCPE-1 and Rab11a/25 than previooulsy anticipated. SNX3 however was re-localised by Rab25. The retromer complex can mediate delivery of endocytic cargoes to late endosomes, and interstingly Rab25 is involved in the sorting of activated α5β1 integrin to late endosomes (Dozynkiewicz et al., 2012), perhaps suggesting Rab25 and retromer may act in concert. Further investigation of these new links between Rab-recycling and ESCPE-1/retromer may help to build a more integrated view of how cargoes can be routed along the endosomal pathway.

### Proximity labelling identitfies functionally relevant associations in migrating cells

Rab4, Rab11 and Rab25 have each been implicated in cell migration, in paricular in 3D matrices, by controlling the trafficking of cagoes such as integrins. Because our datasets included integrins, and the cohort of associated poteins was enriched for GO terms such as ‘cell leading edge’ and ‘focal adhesion’, we reasoned that many of the associated proteins could be involved in motility. Rab4 is required for directional lamellipodial migration by trafficking αvβ3 integrin (Christoforides et al., 2012; White et al., 2007), whilst Rab25 expression promotes a different mode of migration characterised by pseudopod extension in 3D matrix (Caswell et al., 2007; Dozynkiewicz et al., 2012). Rab11, through its effector Rab-coupling protein, controls recycling of α5β1 integrin and cell migration/invasion in cells that express mutant p53 or when Rab4/αvβ3 are inhibited (Caswell et al., 2008; Muller et al., 2009), but we have also found that Rab11 is required for efficient migration in the absence of these factors (Gemperle et al., 2021). We therefore investigated the ability of cells to migrate in a 3D cell-derived matrix in the presence or absence of Rab25 expression after knockdown of candidate machinery closely linked to Rab4a/11a/25 through our experiments, reasoning that Rab25 interacting proteins could be required for migration in the context of Rab25 expression alone, and that the Rab4 related machinery may become redundant when Rab25 is expressed. Indeed, SH3BP5L, a reported Rab25 GEF in vitro which is selectively biotinylated and re-localised by Rab25 had no effect on basal migration of A2780-DNA3 cells, but suppressed migration of A2780-RAB25 cells (Figure 6A, B). Similarly, CLINT1, enriched and more highly biotinylated by Rab4a, is required for migration in 3D-CDM, but only in the absence of Rab25 expression (Figure S6A, B). Interestingly, the selective Rab11 associated protein PI4KB is not required for migration (Figure S6C), which could reflect its role in golgi-mediated processes rather than recycling *per se*.

The dynein adaptor CRACR2A (Rab46) is linked similarly to Rab11 and Rab25 (but not Rab4a) through our proteomic/biotinylation/re-localisation data (Figure 2). Interestingly, knockdown of CRACR2A reduced migration in 3D-CDM in both the presence and absence of Rab25. Our findings further indicate that CRACR2A resides predominatly on a late endosomal/lysosomal compartment. Rab25 endosomes periodically associate with CRACR2A late endosomes, suggesting that cargo exchange could occur in an analogous manner to that seen with Rab25 and CLIC3 in the trafficking of activated α5β1 integrin. How Rab11 might associate with such a pathway, and which cargoes it could deliver is not known, however given that the CLIC3-related CLIC4 is implicated in integrin trafficking and motility it is possible that a parallel pathway handles alternative integrin cargoes in the absence of Rab25.

The finding presented here represent a robust resource cataloguing Rab4a, Rab11a and Rab25 associated proteins that confirm previous findings but also create links to previously unrelated trafficking machineries. Our validation approach, using knock-sideways allows for a functional appraisal of the relevance of candidates that could be more broadly expanded and applied to confirm proximity labelling data which can reveal dynamic interactors. Moreover, linking the robust identification of prey proteins with phenotypes associated with their specific baits has identified new players in the endocytic recycling pathways which regulate cell migration (Figure 7).

## Materials and Methods

### Cell culture

A2780 [DNA3/RAB25] ovarian cancer cell lines (Cheng et al., 2004), were cultured in Roswell Park Memorial Institute-1640 (RPMI-1640) medium (R0883, Sigma). A2780 cells transiently or stably expressing BioID fusion proteins were cultured in RPMI 1640 medium without biotin (R9002-01, US Biological), made up according to the manufacturer’s instructions. Telomerase immortalised fibroblasts (TIFs; (Caswell et al., 2007)) were cultured in Dulbecco’s Modified Eagles Medium (DMEM) (D5796, Sigma). All cell culture medium was supplemented with 10% v/v foetal bovine serum (FBS), 1% antibiotic-antimycotic (Sigma), and 2 mM L-glutamine (Sigma). Cells were maintained at 37°C in a humidified atmosphere with 5% (v/v) CO_2_. Biotin (Sigma) was added to cell culture medium to give the final concentrations indicated for a total of 16 h.

### Antibodies and probes

A range of primary and secondary antibodies were used in this study for immunofluorescence (IF) and western blotting (WB): c-Myc 9E10 (Sigma/M4439; WB 1:1000; IF 1:200) or 9B11 (Cell Signalling/2276S; WB 1:1000; IF 1:100); Rab11 (Invitrogen/715300; WB 1:1000; IF 1:200) α-tubulin DM1A (Abcam/ab7291; WB 1:5000) or YL1/2 (Abcam/ab6160; WB 1:1000); CLINT1 (Abcam/ab223088; WB 1:500); CLINT1 (Thermo Scientific/PA5-60308; IF 1:200); CRACR2A (Proteintech/15206-1-AP; WB 1:1000; IF 1:200); PI4KB (Proteintech/13247-1-AP; WB 1:500); SH3BP5L (Atlas Antibodies/HPA038068; WB 1:500; IF 1:200); RFP (5F8; Chromotek/5f8-100; IF 1:200); Rab11FIP5 (Proteintech/14594-1-AP; IF 1:1000). Streptavidin DyLight®-800 (1:5000) (Thermo Fisher Scientific) was used for western blotting, and Streptavidin-Cy3 (1:200) (Invitrogen) was used for immunofluorescence microscopy. LysoTracker™ Deep Red (1:500000) (Molecular Probes, Invitrogen) and MitoTracker™ Deep Red FM (1:4000-1:8000) (Molecular Probes, Invitrogen) dyes were used for cell imaging.

### Generation of stable cell lines

The lentiviral vector, pCDH tagBFP T2A myc-BirA*-Bax, used as a base for the BioID lentiviral vectors generated for this study, was a gift from Andrew Gilmore (University of Manchester). Rab4a, Rab11a and Rab25 were expanded by polymerase chain reaction (PCR) using primers to introduce XhoI/SalI restriction sites, and ligated to replace myc-BirA*-Bax and generate constructs as detailed in Figure S1A (https://doi.org/10.48420/19391300). Lentiviral particles were produced in HEK293T cells, via a polyethylenimine (PEI)-mediated transfection with pCDH plasmid, pM2G and psPAX2 for 72 h before medium containing viral particles was collected and subjected to centrifugation for 4 min at 1000 rpm and filtration through a 0.45 μm filter. A2780 cells were resuspended in the filtered virus-containing media before plating, and the virus was left on cells for 24 h, after which it was removed and replaced with normal growth media.

TagBFP expressing cells were selected via fluorescence-activated cell sorting (FACS). A2780 cells were resuspended in Ham’s F12 medium (Sigma) supplemented with 25 mM HEPES (Sigma) and 1% antibiotic-antimycotic (Sigma), and filtered through a 50 μm filter. Cells were sorted using an LSR Fortessa cell analyser and BD FACSDiva 8 software (BD Biosciences) based on tagBFP expression.

### Proximity labelling

Cells expressing BioID fusion protein constructs were plated onto tissue culture plates as at a density to ensure near-confluency the following day. More than 4 h after cells were plated biotin (Sigma) was added to cell culture medium (1 μM), and cells were incubated in the presence of biotin for 16 h. BioID cell lysis was carried out at room temperature using a protocol adapted from (Roux et al., 2012). Briefly, cells were PBS washed before addition of the BioID lysis buffer (50 mM Tris pH 7.4, 500 mM NaCl, 0.4% SDS, 5 mM EDTA, 1 mM DTT, supplemented with the protease inhibitors; 100 μg/ml leupeptin (Sigma), 100 μg/ml aprotinin (Sigma), 0.5 mM AEBSF (Calbiochem), 500 μM ALLN (Calbiochem) and 50 μM PD150606 (Calbiochem). Cell lysates were collected using a cell scraper and subjected to needle lysis. Triton X-100 (Sigma) was added to the cell lysates to a final concentration of 2% before needle lysis and addition of an equal volume of 50 mM Tris pH 7.4. Lysates were clarified by centrifugation (16,000xg for 10 min at 4°C). Cell lysates were incubated with 15 μl of MagReSyn® Streptavidin microspheres (per 10 cm plate; ReSyn Biosciences), overnight at 4°C with rotation. Beads were washed twice in wash buffer 1 (2% SDS), once in wash buffer 2 (0.1% deoxycholate, 1% Triton X-100, 500 mM NaCl, 1 mM EDTA, 50 mM HEPES pH 7.4) and once in wash buffer 3 (0.5% NP-40, 0.5% deoxycholate, 1 mM EDTA, 10 mM Tris pH 8.1). Bound proteins were eluted by the addition of 40 μl of 2x sample buffer saturated with biotin (250 mM Tris-HCl pH 6.8, 2% SDS, 10% glycerol, 0.2% Bromophenol blue, 20mM DTT, with >1mM biotin) at 70 °C for 5 min. 10-25% of eluted samples were retained for western blot analysis and the remaining sample was used for mass spectrometry analysis.

### Mass spectrometry sample preparation

Eluted proteins from BioID experiments were subjected to SDS-PAGE at 100 V for 8 min. Gels were stained with InstantBlue™ Protein Stain (Expedeon) for 10 min and subsequently washed in ddH2O at 4°C overnight with gentle shaking. ‘Gel-top’ protein bands were excised and transferred to a 96-well perforated plate containing ddH2O and then centrifuged to remove liquid. Gel pieces were de-stained sequentially using 50% acetonitrile (ACN)/50% 25 mM NH4HCO3, followed by 100% ACN. De-stained gel pieces were dried using a vacuum centrifuge and proteins reduced (10 mM DTT in 25 mM NH4HCO3) and then alkylated (55 mM iodoacetamide/25 mM NH4HCO3) in the absence of light. Gel pieces were washed alternately in 25 mM NH4HCO3 then 100% ACN and dried using a vacuum centrifuge and trypsin (1.25 ng/μl in 25 mM NH4HCO3) digested overnight. Peptides were collected in 96-well collection plates by centrifugation washed in 99.8% ACN, 0.2% formic acid (FA) then 50% ACN, 0.1% FA. Peptide solutions were dried to completion using a vacuum centrifuge and resuspended in 5% ACN/0.1% FA. Desalting of peptides was carried out using POROS Oligo R3 beads (ThermoFisher). Beads were washed with 50% CAN, then 0.1% FA, and peptides eluted twice with 50% ACN/0.1% FA. Peptide solutions were dried to completion using a vacuum centrifuge and resuspended in 20 μl 5% ACN, 0.1% FA for mass spectrometry analysis.

### Mass spectrometry analysis

Peptides were analysed using liquid chromatography tandem mass spectrometry (LC-MS/MS) using an Orbitrap Elite™ (Thermo Fisher) in the Biological Mass Spectrometry core facility, University of Manchester. Peptides were automatically selected for fragmentation by data-dependent analysis. Four biological replicates of each sample were analysed in one batch on the same day in the following order myc-BirA*-Rab25, myc-BirA*-Rab4a, myc-BirA*-Rab11a, myc-BirA* only, with all biological replicates of each bait analysed consecutively before replicates of the next bait were analysed. All samples were analysed using 2 h runs and LC column equilibrated with a pooled sample (consisting of an equal mix of all samples) between each set of baits.

For MS1 intensity-based analysis, data were analysed using MaxQuant (version 1.6.2.10). Default MaxQuant parameters were used, with fixed modifications of cysteine carbamidomethylation and variable modifications of methionine oxidation, protein N-terminus acetylation and lysine biotinylation. Label-free quantification (LFQ) was selected and match between runs was turned on. Data were searched against a human SwissProt/UniProt database (downloaded March 2018). LFQ intensities from MaxQuant analysis were analysed by SAINTexpress (version 3.6.3), using default parameters.

A range of bioinformatic analyses were performed on the mass spectrometry data. PPI network visualisation was performed using Cytoscape, with proteins mapped onto the Homo sapiens Biological General Repository for Interaction Dataset (BioGRID) 3.4.162 database. Proportional Venn diagrams were generated using an online tool (http://eulerr.co/). Dot plots of the high-confidence proximal proteins identified to each Rab GTPase were generated by ProHits-viz (Knight et al., 2017) (https://prohits-viz.lunenfeld.ca/) from SAINTexpress output data. Gene ontology analysis was performed using clusterProfiler (Yu et al., 2012) in R. Principal component analysis (using prcompf base package), pairwise comparisons and the visualisation of biotinylated peptides were all performed in R and plotted using ggplot2.

### Immunofluorescence

For staining with anti-myc (9E10) antibody, cells were methanol fixed in −20°C methanol for 10 min at −20°C followed by −20°C acetone for 1 min at room temperature. For all other antibody staining procedures, cells were fixed in 4% (w/v) paraformaldehyde (Sigma) for 15 min at room temperature, before permeabilisation with 0.2% Triton-X 100 (Sigma) in PBS- for 5 min at room temperature.

Fixed cells were blocked in 1% (w/v) heat-denatured BSA (Sigma) in PBS- with 10% (v/v) FBS and incubated with primary/secondary antibodies diluted in 1% heat-denatured BSA in PBS- for 1 h at room temperature. Cells were washed five times with PBS and once with ddH2O before mounting with either ProLong™ Gold Antifade Mountant (Molecular Probes, Invitrogen) or ProLong™ Diamond Antifade Mountant (Molecular Probes, Invitrogen). Images were acquired using a Leica TCS SP8 AOBS inverted confocal microscope using a 63x APO objective, using hybrid detectors with the appropriate detection mirror settings and sequential image collection, capturing z-stacks using LAS X software (Leica).

### Knock sideways imaging and analysis

Cells were nucleofected (VCA-1002, Lonza) with required amount of plasmid DNA using programme A-23 (Lonza). For live cell experiments the following amounts of plasmid DNA were used; 1 μg Mito-mCh(K70N)-FRB, 3 μg mCh-Rab11FIP5 (gift from Gift from Andrew Lindsay, University College Cork) and 1 μg of the required GFP-FKBP, to achieve optimal protein expression. Rab11a/Rab4/Rab25 was introduced to GFP-FKBP-C1 plasmid (gift from Steve Royle, University of Warwick) by PCR.

Cells were nucleofected as indicated and imaged after 24 h. Cell growth medium was replaced with Opti-Klear™ Live Cell Imaging Buffer (Tebu-bio) supplemented with 10% FBS and MitoTracker™ Deep Red FM (1:8000) for at least 30 min before image acquisition. Cells were imaged using 3i spinning disk confocal microscopy. Single-section confocal images were collected using a CSU-X1 spinning disc confocal (Yokagowa) on a Zeiss Axio-Observer Z1 microscope with a 63x/1.46 α Plan-Apochromat objective, a Prime 95B Scientific CMOS camera (Photometrics) and motorised XYZ stage (ASI). Lasers were controlled using an AOTF through the laser stack (Intelligent Imaging Innovations, 3i) allowing both rapid ‘shuttering’ of the laser and attenuation of the laser power. Slidebook software (3i) was used to capture images of approximately 10 positions per condition, every 5 min for 1 h. Rapamycin (Sigma) to a final concentration of 200 nM was added to cells on the microscope immediately after the first time point image was acquired to stimulate the rerouting of GFP-FKBP constructs to the mitochondria.

For the knock sideways fixed approach, we simplified nucleofection by using 2 μg pMito-iRFP670-FRB (Gemperle et al., 2021) with 1μg of the required GFP-FKBP. The following day rapamycin was added to the cell growth medium at a final concentration of 200 nM for 4 h. Cells were then fixed and stained for the relevant candidate protein (described above), detected by an Cy-3 secondary antibody, using Decon Vision deconvolution microscopy. Images were acquired using an Olympus IX83 inverted microscope using Lumencor LED excitation, with a 60x/ 1.42 Plan Apo objective and a Retiga R6 (Q-Imaging) CCD camera. Metamorph software (Molecular Devices) was used to capture z-stacks, and these were then deconvolved using the Huygens Pro software (SVI) with default settings.

All images were processed using ImageJ, relocalisation of GFP-FKBP-(Rabs) and mCh-Rab11FIP5 to mitochondria was analysed using the Colocalization finder plugin and scored based on Pearson’s correlation coefficient (no thresholding, only noise subtraction in ScatterPlot). KnockSideways relocalisation dynamics was corrected for signal crosstalk, normalized to scale 0-1 and fitted with non-linear fit using GraphPad Prism and T_1/2_ determined as time necessary to relocalise 50 % of total GFP signal (equilibrium) to Mitochondria. For fixed deconvoluted images, a mitochondrial mask was created using a custom made script and the mean pixel intensity of candidate protein was compared between mitochondria and the cell body (https://doi.org/10.48420/19391597)

### 3D-cell derived matrix (CDM) migration analysis

Lipofectamine transfection was used to introduce siRNAs (Qiagen) into A2780 cells for protein depletion: CLINT1 siRNA_1: 5’ CAGGCTTCGTGAAGAGCGAAA (SI04178748, Hs_CLINT1_1); CLINT1 siRNA_3, 5’ ATGGTAAGGATCAAGGTATAA (SI04230926, Hs_CLINT1_3); CRACR2A siRNA_1 5’ TACCGTGTGACGGAGAGTCTA (SI04218130, Hs_EFCAB4A_1); CRACR2A siRNA_4 5’ AAGAAGGAGGAACCTCATTTA (SI04322682, Hs_EFCAB4B_4); PI4KB siRNA_5 5’ TCGGCTGATAGTGGCATGATT (SI00605850, Hs_PIK4CB_5); PI4KB siRNA_6 5’ CGACATGTTCAACTACTATAA (SI02660077, Hs_PIK4CB_6); SH3BP5L siRNA_2 5’ ATGCACAACGCTGCTCGAGAA (SI04172315, Hs_SH3BP5L_2); SH3BP5L siRNA_3 5’ CACGTCAGTCTGGACGGCCAA (SI04192993, Hs_SH3BP5L_3).

CDMs were prepared as described previously (Caswell et al., 2007; Cukierman et al., 2001), on either tissue-culture plastic multi-well plates (for long-term time-lapse microscopy) or 35 mm glass-bottomed dishes (MatTek) (for live imaging). Briefly plates were coated with 0.2% gelatin (Sigma) in PBS before cross-linking with 1% glutaraldehyde (Sigma) in PBS. Plates were PBS washed and quenched with 1 M glycine (Fisher Scientific) in PBS before washing and equilibration in DMEM. TIFs were plated onto the prepared plates at a density to ensure confluency the following day. TIFs were grown for 8–10 days, with the medium changed for DMEM supplemented with 25 μg/ml ascorbic acid (Sigma) 24 h after seeding and every 48 h after that. Matrices were denuded of cells with extraction buffer (20 mM NH4OH, 0.5% (v/v) Triton X-100, in PBS) and washed twice with PBS+ (Dulbecco’s Phosphate Buffered Saline with calcium chloride and magnesium chloride, Sigma), incubated with 10 μg/ml DNase I (Roche) and washed three times with PBS+ before A2780 cells were plated at a density of 4×104 cells per well of 6 well plate or 35 mm dish. Cells were allowed to spread and start migrating for at least 4 h before use in experiments.

Images were acquired on an Eclipse Ti inverted microscope (Nikon) using either a 20x/0.45 SPlan Fluor or a 10x/0.3 Plan Fluor objective, and a pE-100 LED (CoolLED) light source. Images were collected using a Retiga R6 (Q-Imaging) camera, and cells were maintained at 37°C and 5% CO2 for the duration of imaging. NIS Elements AR.46.00.0 software was used to acquire images of six positions per well, every 10 min for 16 h. For cell speed analysis, at least 5 cells were individually manually tracked per position (giving a minimum of 30 cells tracked per condition per repeat, at least 90 cells in total) using the ImageJ MTrackJ plugin, and this was used to calculate the average cell speed (Meijering et al., 2012).

### Live cell imaging

Cells were nucleofected with mCherry-CRACR2A-L (Addgene #79593) and EGFP-Rab25 (Caswell et al., 2007)as indicated and seeded onto 35 mm glass-bottomed CDMs 24 hours later. Cell growth medium was replaced with Opti-Klear™ Live Cell Imaging Buffer (Tebu-bio) supplemented with 10% FBS at least 30 min before image acquisition. Cells were incubated with LysoTracker™ Deep Red (1:500000) diluted in Opti-Klear for at least 30 min before image acquisition and maintained at 37°C during imaging.

Live-cell imaging was carried out using spinning disk confocal microscopy (3i Marianis). Single-plane images were collected CSU-X1 spinning disc confocal (Yokagowa) on a Zeiss Axio-Observer Z1 microscope with a 63x/1.46 α Plan-Apochromat objective and a Prime 95B Scientific CMOS camera (Photometrics). Lasers were controlled using an AOTF through the laser stack (Intelligent Imaging Innovations, 3i) allowing both rapid ‘shuttering’ of the laser and attenuation of the laser power. Slidebook software (3i) was used to capture images of cells every 30 s for 10 min. Images were processed and analysed using ImageJ.

### Statistics

Data were tested for normality and one-way ANOVA with Tukey or Holm-Sidak post hoc test used for multiple comparisons as indicated in legend. Where data were not normally distributed, Kruskal-Wallis test with Dunn’s multiple comparison. All statistical analysis has been performed with GraphPad Prism software, where *** denotes p < 0.001, ** denotes p < 0.01, and * denotes p < 0.05. Data represent at least three independent experiments; n numbers and p values are described in relevant figure legends.

### Data Availability

Custom scripts/macro codes are deposited on Figshare (https://doi.org/10.48420/19391597). Annotated plasmid maps generated in this study can be download from https://doi.org/10.48420/19391300. Plasmids can be provided upon reasonable request from corresponding author. The mass spectrometry proteomics data have been deposited to the ProteomeXchange Consortium via the PRIDE (Perez-Riverol et al., 2022) partner repository with the dataset identifier PXD033693.

## Acknowledgements

The Bioimaging Facility microscopes used in this study were purchased with grants from BBSRC, Wellcome and the University of Manchester Strategic Fund. We thank Peter March and Steven Marsden for their help with microscopy, and David Knight and the BioMS facility for assistance with mass spectrometry. The Flow Cytometry Core Facility is supported in part by the University of Manchester with assistance from MRC Grant ref MR/L011840/1. This project received funding from the European Union’s Horizon 2020 research and innovation programme under grant agreement No [836212]; Wellcome Trust studentships (109330/Z/15/A; BW and 220005/Z/19/Z; MH); Worldwide Cancer Research (14-1226); the MRC (MR/R009376/1); Cancer Research UK (DCRPGF\100002; CF is funded by the Cancer Research UK Manchester Centre [C147/A25254] Non-Clinical Training Programme and the Wellcome Trust Centre for Cell Matrix research is funded by grant 203128/A/16/Z.

---

**Supplementary Figure 1:**
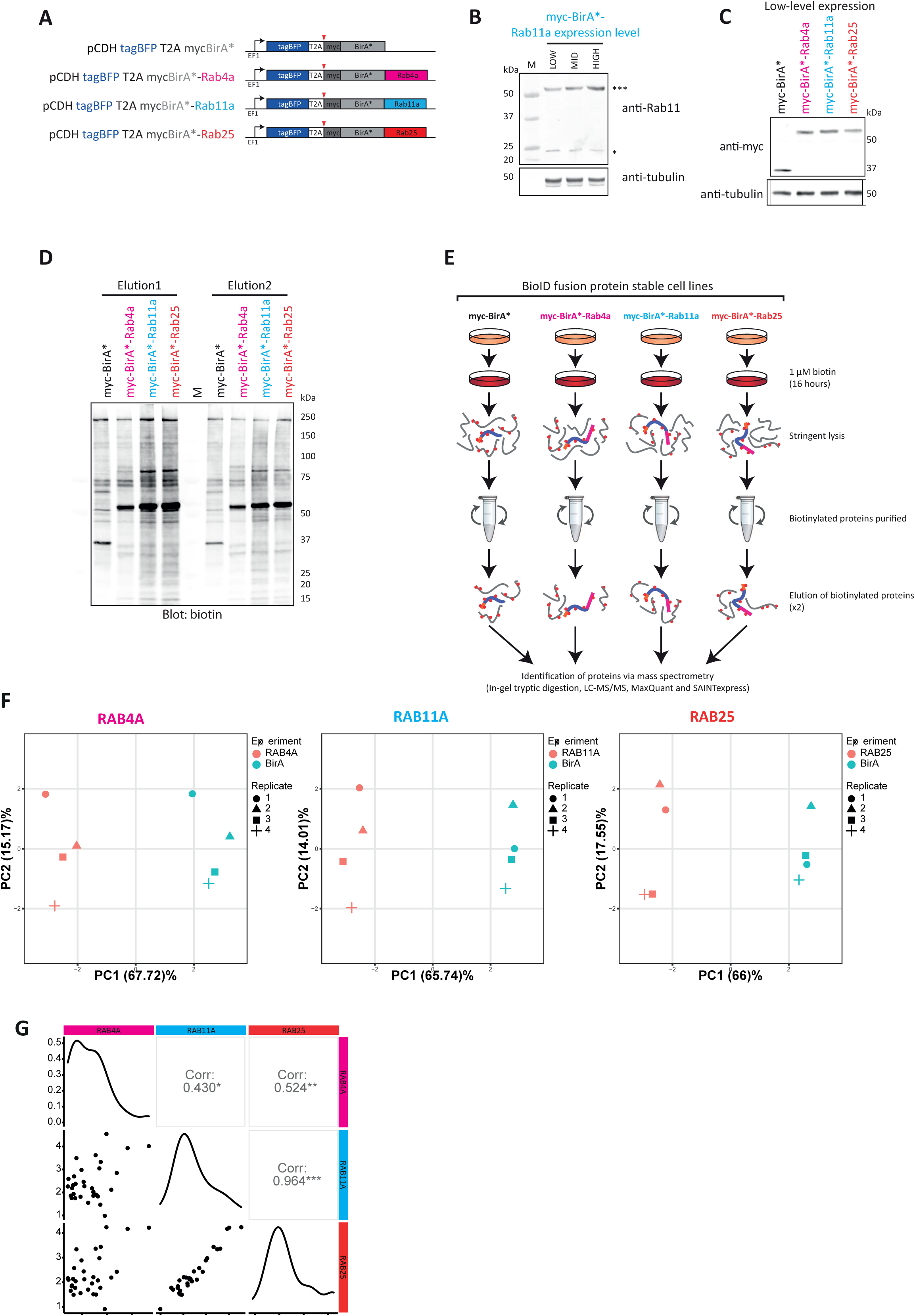
Generation of a high-quality library of Rab4a/11a/25 associated proteins using proximity biotinylation. **(A)**. Schematic of the lentiviral BioID constructs used to generate stable cell lines. Co-expression occurs before co-translational cleavage of the T2A peptide, enabling the separate expression of tagBFP and the fusion protein. **(B)**. Expression of myc-BirA*-Rab11a in stable cell lines was confirmed by western blotting using a Rab11 antibody (upper panel), and α-tubulin as a loading control (lower panel). *, endogenous Rab11; ***, myc-BirA*-Rab11a. Molecular weight marker, M, is shown in kDa. **(C)**. Expression of BirA* fusion proteins in stable cell lines was confirmed by western blotting using a myc antibody (myc, upper panel), and α-tubulin as a loading control (lower panel). **(D)**. A2780 cells stably expressing BirA* fusion proteins were plated for 7-8 h and supplemental biotin (1μM) added for 16 h as indicated. Cells were lysed and biotinylated proteins were affinity purified using MagReSyn® streptavidin microspheres, with two subsequent elutions carried out in 2x sample buffer saturated with biotin, for 5 min at 70 °C. A proportion of resultant samples (10%) were analysed by western blot, and a representative blot from one of the four replicates is shown. Biotinylated proteins detected by Alexa800-streptavidin (biotin). Molecular weight marker, M, is shown in kDa. **(E)**. Experimental pipeline. Identification of enriched proteins was carried out by LC-MS/MS, followed by analysis by MaxQuant and SAINTexpress. **(F)**. Principle component analysis of label-free quantification intensities (from MaxQuant) for the identified affinity-purified proteins. The analyses were carried out on all replicates of the BirA*-only control and all replicates of each BirA*-Rab GTPase fusion protein in turn. **(G)**. Pairwise comparison of the Rab4a, Rab11a and Rab25 fold changes (log_2_FoldChange) over myc-BirA* of the 31 high-confidence proximal proteins shared by all 3 Rab GTPases, Pearson correlation values are shown.

**Supplementary Figure 2:**
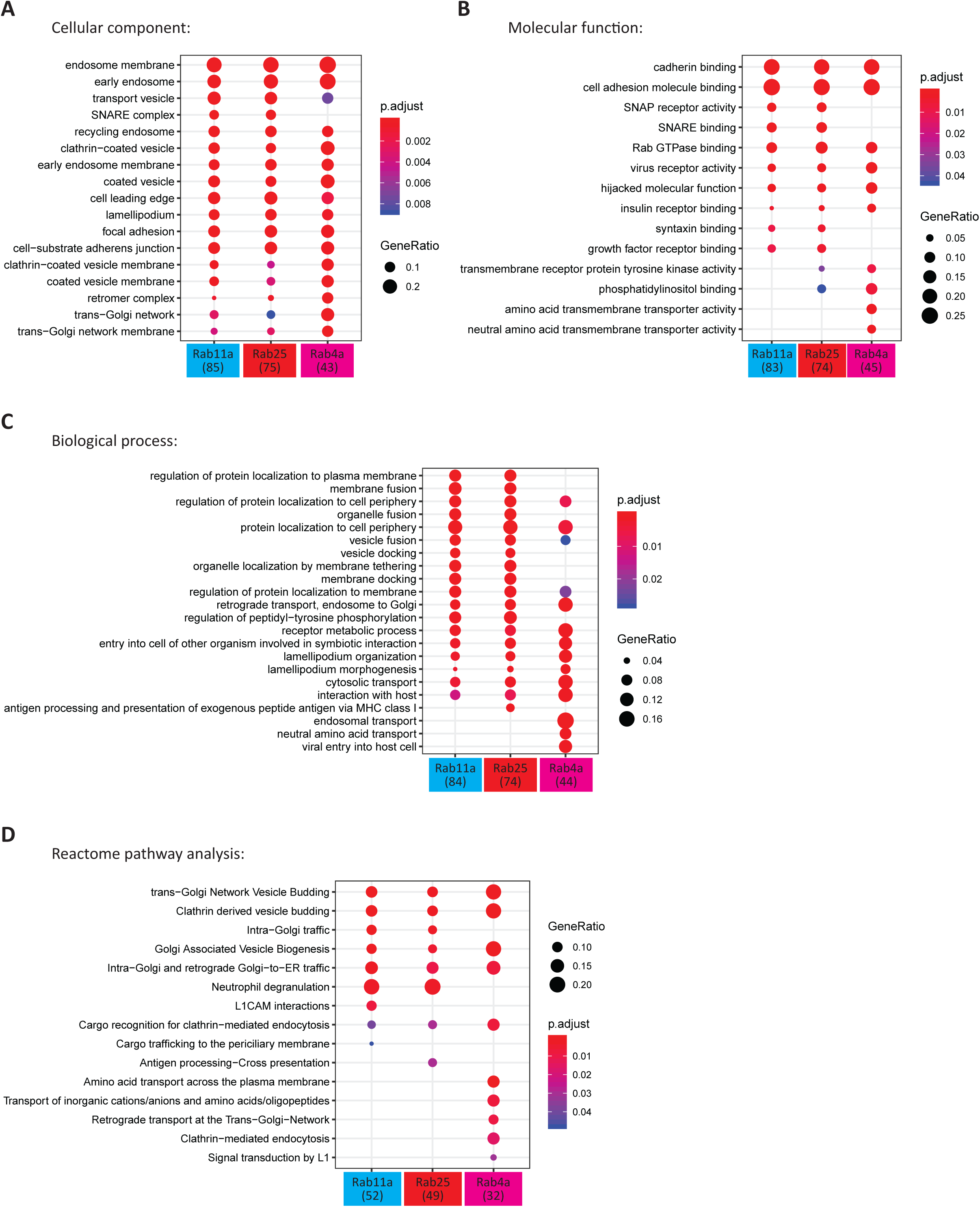
Gene ontology analysis of high-confidence Rab GTPase proximal proteins. GO analysis of the high-confidence proximal proteins for Rab4a, Rab11a and Rab25 was carried out. The top 10 enriched GO terms under the stated categories are displayed for each bait (p-value ≤ 0.05), nodes coloured according to p.adjust, adjusted p-value. Number of proteins recognised per bait in brackets; GeneRatio, proportion of total proteins identified in each GO term. (A) Cellular component GO category (B) Molecular function GO category (C) Biological process GO category (D) Reactome pathway analysis terms.

**Supplementary Figure 3:**
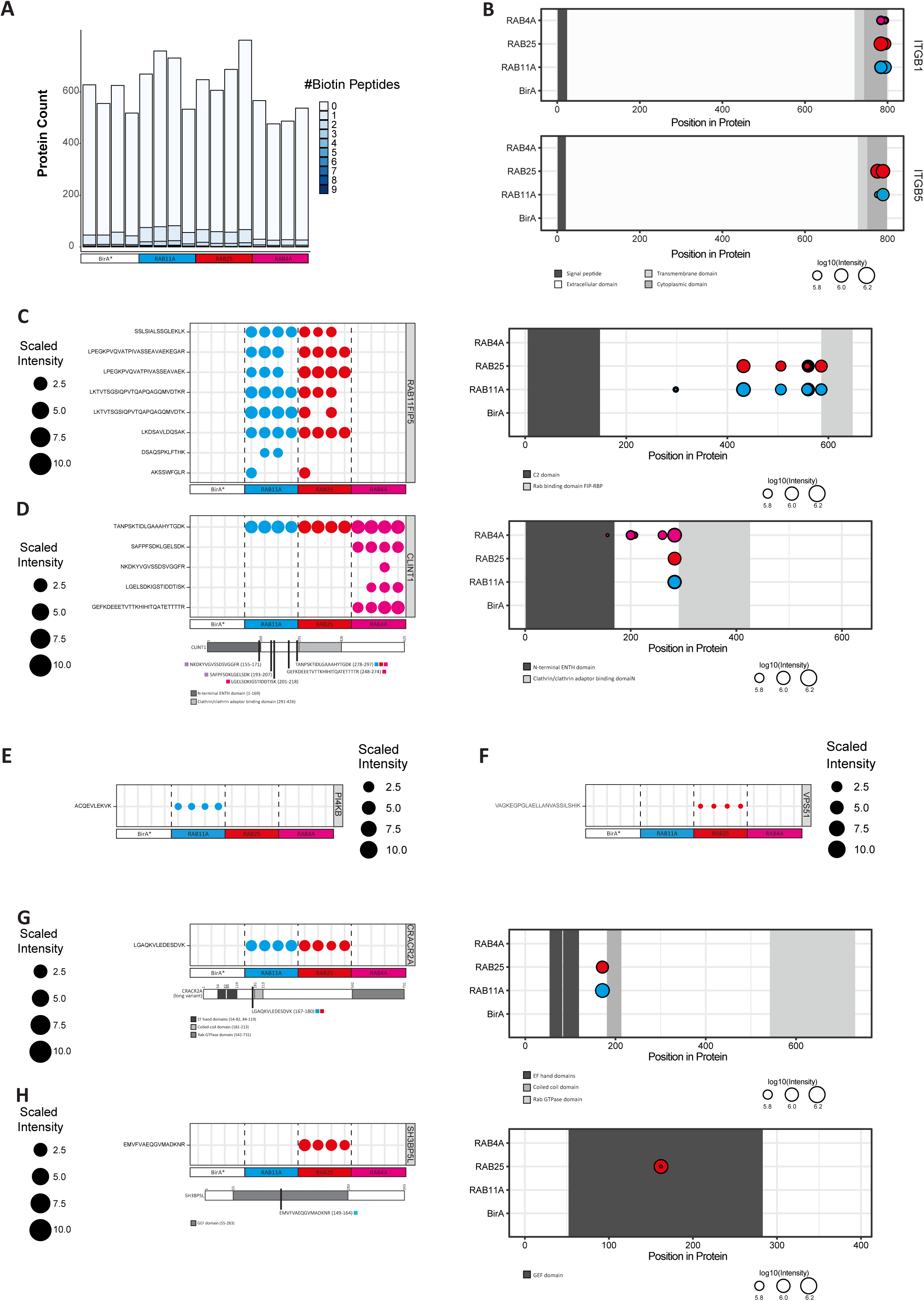
Mapping biotin modifications in prey proteins. **(A)**. Numbers of biotinylated peptides identified for each protein group in each replicate, colour coded according to the number of biotinylated peptides identified for each. **(B-H)**. Biotin modifications within prey proteins were mapped according to peptide and/or position in the prey protein.

**Supplementary Figure 4:**
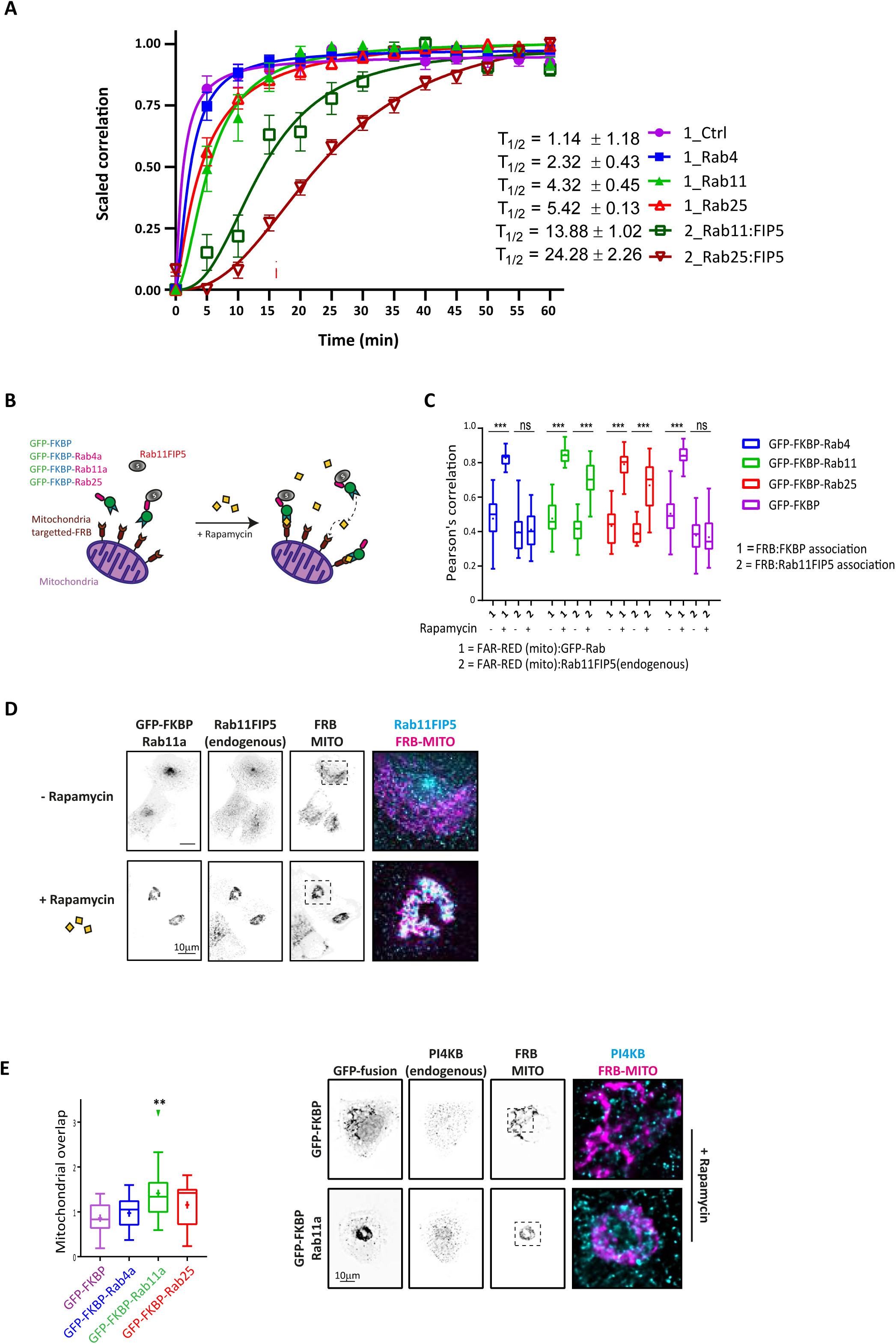
Knock-sideways validation of proximity labelling-identified preys. **(A)**. Quantification of the redistribution of GFP-tagged proteins and corresponding localisation of mCherry-Rab11-FIP5 in live A2780 cells expressing iRFP670-FRB and GFP-FKBP or GFP-FKBP-Rab4a/11a/25 and mCherry-Rab11-FIP5 (for images see Figure 4C, see methods). **(B)**. Schematic illustration of knock-sideways, where mitochondria-targeted FRB/rapamycin is used to induce the re-localisation of GFP-FKBP-tagged Rabs and associated protein complexes. A2780 cells expressing iRFP670-FRB and GFP-FKBP-Rabs treated +/− rapamycin (200 nM 4h) were fixed and stained for endogenous Rab11-FIP5 **(C, D)** or PI4KB **(E)**. Re-distribution of GFP-FKBP fusion (1) and mCherry-Rab11-FIP5 (2) were analysed by Pearson’s correlation (B; at least 19 cells/condition; statistical analysis with ANOVA/Holm-Sidak post hoc test; ***p<0.001), representative images from at least 3 independent experiments are show in (C; scale bar=10μm). Redistribution of PI4KB was analysed by quantifying mitochondrial overlap (D; at least 11 cells/condition; statistical analysis with ANOVA/ Holm-Sidak post-hoc test; **p<0.01) representative images from at least 3 independent experiments are shown (scale bar=10μm).

**Supplementary Figure 5:**
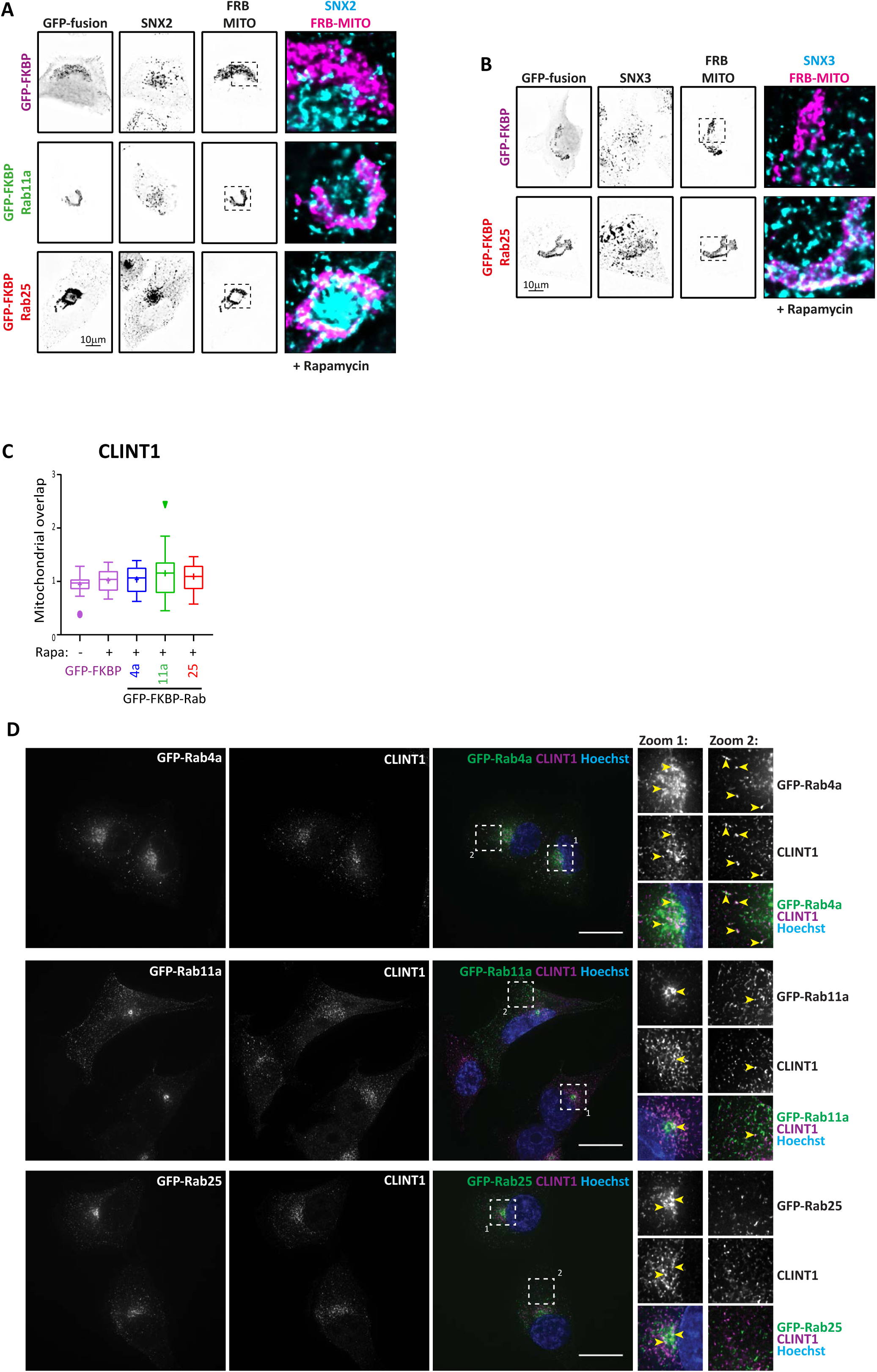
Sorting nexins are selectively recruited to Rab11a and Rab25. A2780 cells expressing iRFP670-FRB and GFP-FKBP-Rabs treated with rapamycin (200 nM 4h) were fixed and stained for endogenous SNX2 (A) or SNX3 (B). Representative images from at least 3 independent experiments are shown (scale bar=10μm). (C). A2780 cells expressing iRFP670-FRB and GFP-FKBP-Rabs treated +/− rapamycin (200 nM 4h) were fixed and stained for endogenous CLINT1. Redistribution of CLINT1 was analysed by quantifying mitochondrial overlap (see methods; at least 13 cells/condition; statistical analysis with ANOVA/Tukey post-hoc test;). (D) A2780 cells expressing GFP-Rabs were fixed and stained for CLINT1, and imaged by deconvolution microscopy. Representative images from at least 3 independent experiments are show in (C; scale bar=20μm).

**Supplementary Figure 6:**
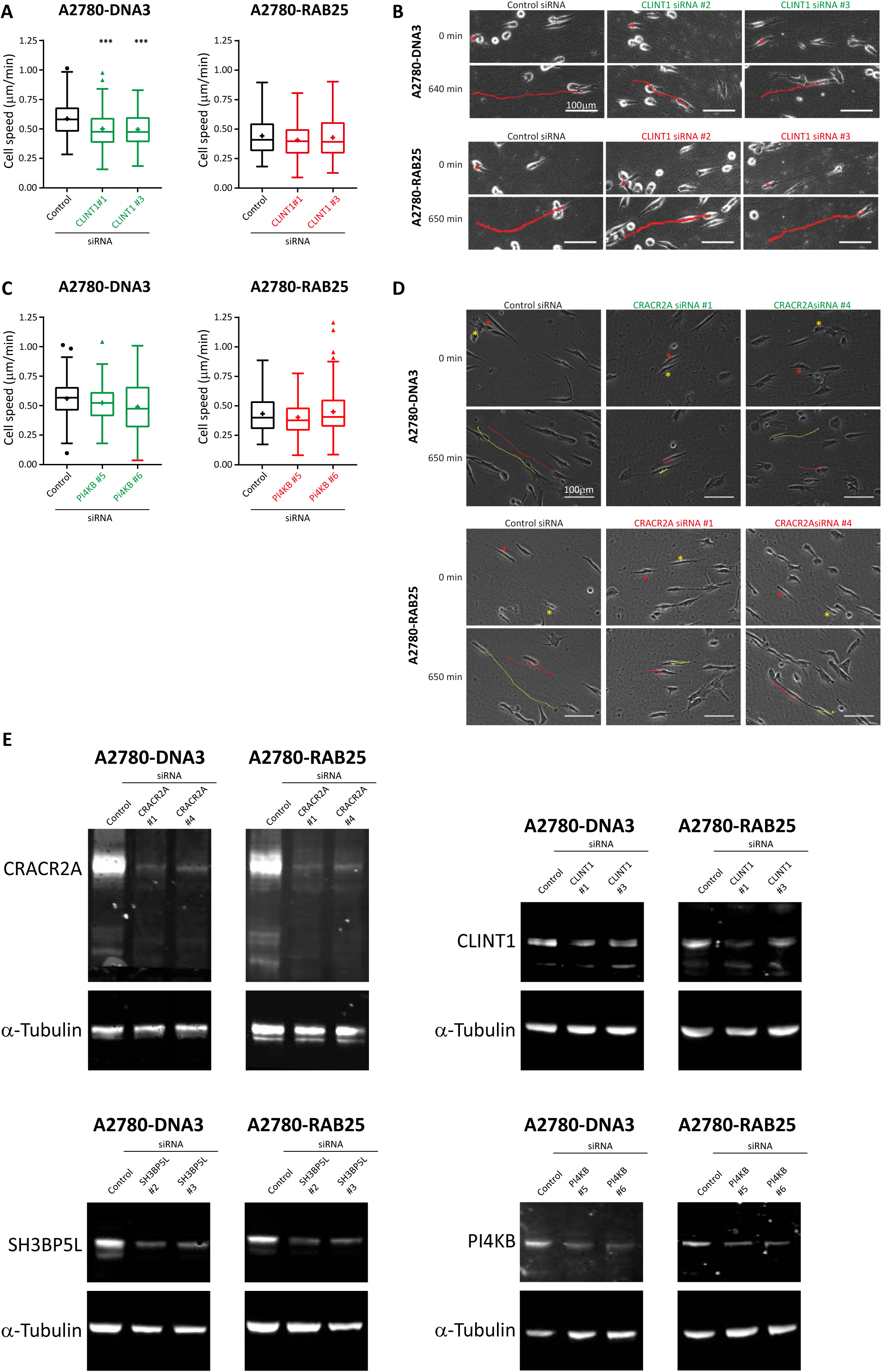
Recycling machineries required for cell migration in 3D-matrix. (A-D). A2780-DNA3 and A2780-Rab25 cells were depleted of CLINT1 or PI4KB by siRNA and seeded into cell-derived matrix for 4 hours before migration was analysed by brightfield timelapse imaging for >16 hours. Cell speed was analysed by manual tracking (A, C; n≥90 cells from 3 independent experiments; statistical analysis with Kruskal-Wallis/Dunn’s multiple comparisons test; ***p<0.001; note: A2780-Rab25 experiments performed simultaneously); representative images from at least 3 independent experiments are shown (B, D; scale bar=100μm). (E). A2780-DNA3 and A2780-Rab25 cells were depleted of CRACR2A, CLINT1, SH3BP5L or PI4KB by siRNA and knockdown analysed by western blot after 48 hours (corresponding to the endpoint of migration analysis). Images are representative of 3 independent experiments.

**Supplementary Table 1: High confidence Rab proximal proteins identified by BioID**

SAINTexpress output for the statistical significance of prey proteins identified by BioID for RAB4A, RAB11A and RAB25.

**Supplementary Table 2: Longlist Rab proximal proteins identified by BioID**

Proteins enriched 2-fold in at least 3 of 4 repeats compared to BirA control. For these proteins the number of unique peptides identified, LFQ intensities in each of four BioID experiments, average LFQ (null values in BirA control were pseudocounted as 0.00001 to give an enrichment ratio) and Rab/BirA enrichment ratio are shown

**Supplementary Table 3: Peptide modifications identified (including biotinylation)**.

All biotinylated peptides identified by BioID, in each sample, for RAB4A, RAB11A and RAB25.

